# Contractile ring constriction and septation in fission yeast are integrated mutually stabilizing processes

**DOI:** 10.1101/2021.06.25.449700

**Authors:** Sathish Thiyagarajan, Zachary McDargh, Shuyuan Wang, Ben O’Shaughnessy

## Abstract

In common with other cellular machineries, the actomyosin contractile ring that divides cells during cytokinesis does not operate in isolation. Contractile rings in animal cells interact with contiguous actomyosin cortex, while ring constriction in many cell-walled organisms couples tightly to cell wall growth. In fission yeast, a septum grows in the wake of the constricting ring, ensuring cytokinesis leaves two daughter cells fully enclosed by cell wall. Here we mathematical modeled the integrated constriction-septation system in fission yeast, with a kinetic growth model evolving the 3D septum shape coupled to a molecularly explicit simulation of the contractile ring highly constrained by experimental data. Simulations revealed influences in both directions, stabilizing the ring-septum system as a whole. By providing a smooth circular anchoring surface for the ring, the inner septum leading edge stabilized ring organization and tension production; by mechanically regulating septum circularity and in-plane growth, ring tension stabilized septum growth and shape. Genetic or pharmacological perturbation of either subsystem destabilized this delicate balance, precipitating uncontrolled positive feedback with disastrous morphological and functional consequences. Thus, high curvature septum irregularities triggered bridging instabilities, in which contractile ring segments became unanchored. Bridging abolished the local tension-mediated septum shape regulation, exacerbating the irregularity in a mutually destabilizing runaway process. Our model explains a number of previously mysterious experimental observations, including unanchoring of ring segments observed in cells with mutations in the septum-growing β-glucan synthases, and irregular septa in cells with mutations in the contractile ring myosin-II Myo2. Thus, the contractile ring and cell wall growth cellular machineries operate as a single integrated system, whose stability relies on mutual regulation by the two subsystems.

## Introduction

Cellular machineries do not operate in isolation, and a full understanding of their mechanisms should embrace coupled systems. The contractile ring is coupled to the contiguous cortical cytoskeleton in animal cells (Reymann, Staniscia, Erzberger, Salbreux, & Grill, 2016; Sedzinski et al., 2011; Turlier, Audoly, Prost, & Joanny, 2014), to cell wall synthesis in fungi and other cell-walled organisms (Fang et al., 2010; Ramos et al., 2019; Thiyagarajan, Munteanu, Arasada, Pollard, & O’Shaughnessy, 2015), and generally to membrane delivery to the furrow (VerPlank & Li, 2005; N. Wang, Lee, Rask, & Wu, 2016).

For cell-wall enclosed organisms, growth of new cell wall material is essential during the physical division of the cell, so that the two daughter cells can be furnished with functional cell wall. In fission yeast, synthesis of the septum, or new cell wall, is performed by membrane-bound β-glucan synthases Bgs1, Bgs3, Bgs4 and the α-glucan synthase Ags1 in the wake of the constricting ring, Figure 1 (Cortes et al., 2005; Cortes, Ishiguro, Duran, & Ribas, 2002; Konomi, Fujimoto, Toda, & Osumi, 2003; Martin, Garcia, Carnero, Duran, & Sanchez, 2003). The septum is a three-layered cylindrical annulus with a tapering cross-section of mean thickness ~ 200 nm, Figure 2A (Cortes et al., 2007; Garcia Cortes, Ramos, Osumi, Perez, & Ribas, 2016). It joins the cell wall proper at its outer boundary and the ring lines its innermost surface.

**Figure 1.**
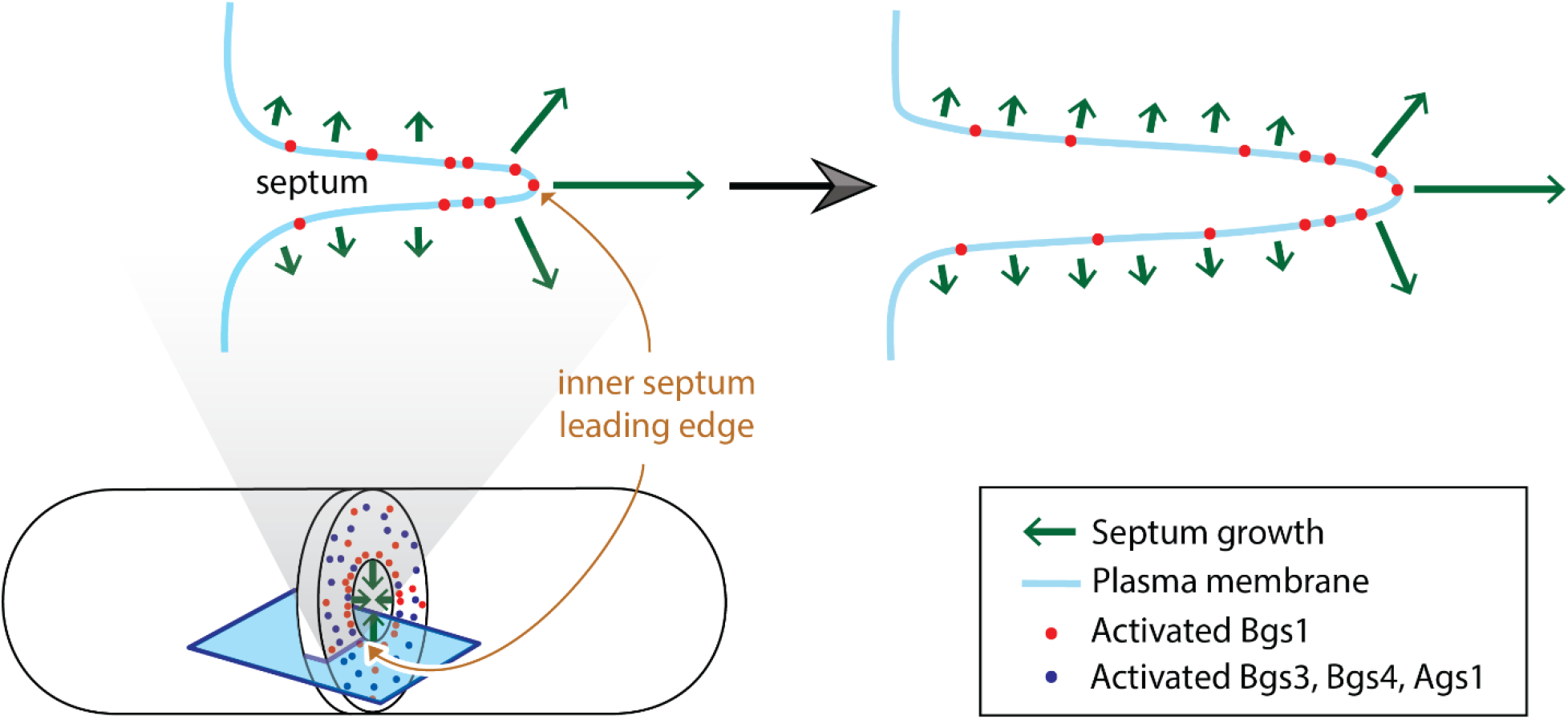
Synthesis of new cell wall material by membrane-bound active glucan synthases grow the septum inward in fission yeast Bottom: schematic of a dividing fission yeast cell illustrating septum growth and glucan synthase distribution at the division plane. Cross section of the septum along the blue plane shown at top. Top: schematic of the septum cross-section showing inward movement of the inner septum leading edge and surrounding regions due to growth produced by membrane-bound glucan synthases. Only activated Bgs1 are shown, as these are the principal synthesizers of septum that evolve the shape. Septum growth by synthases at the inner leading edge closes down the septum hole by advancing the edge. Growth also thickens the septum but at a much lower rate due to activated synthases at the rear. The difference in rates is due to differences in glucan synthase distribution. The density of the β-glucan synthase Bgs1 that primarily synthesizes the fast-growing primary septum is highest at the membrane adjacent to the ring and tapers off further away from the ring (G. Cortés et al., 2015; Goss, Kim, Bledsoe, & Pollard, 2014). The other synthases Bgs3, Bgs4, and Ags1 that are primarily responsible for synthesizing the slow-growing secondary septum are distributed throughout the membrane (Cortes et al., 2005; Cortes et al., 2012; Munoz et al., 2013). Schematics are not to scale.

**Figure 2.**
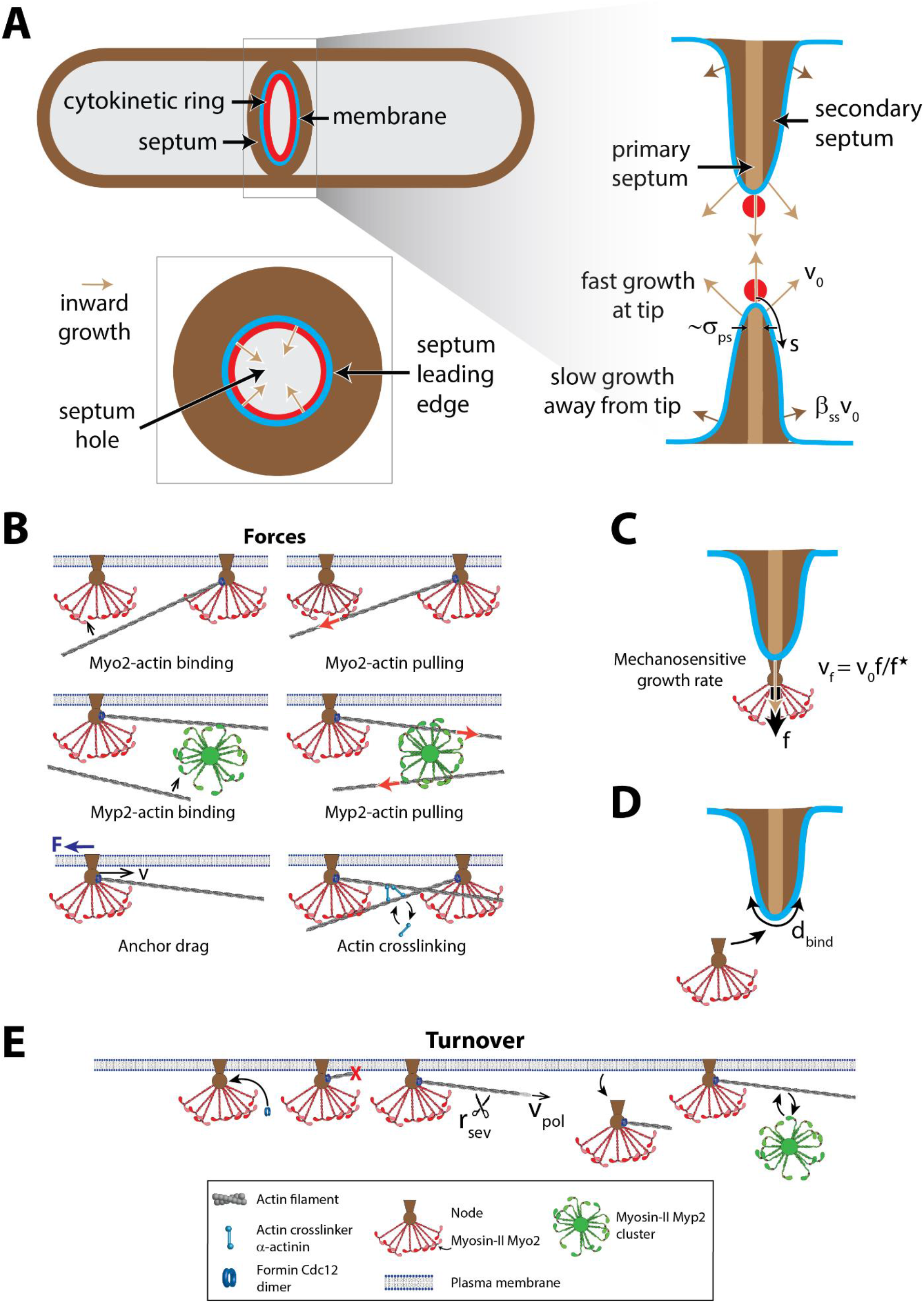
Three-dimensional mathematical model of septum growth coupled to ring constriction. **(A)** Top left: Schematic of a fission yeast cell undergoing division. The coupled processes are constriction of a contractile ring (red) and the growth of new cell wall material or septum in its wake (brown), synthesized by glucan synthases distributed throughout the membrane (blue). Bottom left: Top view of the division plane illustrating the two-dimensional septum hole enclosed by the leading edge of the septum. Right: Side view of the septum, which is a three-layered tapering structure with an inner fast-growing primary septum and two outer layers of slow-growing secondary septum. Distance from the ring is parameterized by *s*. Growth of the primary septum pushes the leading edge inward at a rate υ_0_. Growth rate tapers over a distance *s*~*σ*_ps_, with the secondary septum thickening at the much slower rate of *β*_ss_υ_0_. Schematics are not to scale. **(B)** Schematic of force-producing interactions in the ring. Dimers of the myosin II isoform Myo2 and formin Cdc12 dimers colocalize in membrane-bound nodes. Myo2 and the clusters of the unanchored isoform Myp2 bind actin filaments anchored to the membrane via formins at their barbed ends and exert pulling forces parallel to the filaments which are resisted by membrane drag forces on the node anchors. Crosslinkers bind to filament segments whose separation is within a specified distance. **(C)** Schematic of growth rate acceleration at the leading edge of the septum due to mechanosensitivity of septum growth. An inward force *f* on a node increases growth rate by υ_0_*f*/ *f*\* in its neighborhood of size *b*_node_, where υ_0_ is the characteristic growth rate at the septum leading edge and *f*\* is a critical force. For details, see Materials and Methods. **(D)** Schematic of new nodes binding the septum surface in a zone within a distance *d*_bind_ of the septum leading edge. **(E)** Schematic showing turnover of molecular components in the ring. Formin dimers bind nodes which have bound the septum and synthesize filaments at a rate υ_pol_ away from the membrane. Cofilin-mediated severing occurs at a rate *r*_sev_ with uniform probability per unit length. Nodes and Myp2 clusters unbind from the ring at a characteristic probability per unit time. Both Myo2 and Myp2 clusters bind and unbind actin filaments. **(B)**, **(E)** adapted from (McDargh et al., 2021).

Previous studies suggest tight coupling between the contractile ring and septation in fission yeast. Mutations in ring proteins alter the septum growth rate (Bellingham-Johnstun, Anders, Ravi, Bruinsma, & Laplante, 2021; Sladewski, Previs, & Lord, 2009). When the ring is removed using treatment with the actin monomer sequestering drug Latrunculin A, the septum stops ingressing if the treatment is performed early, but ingresses at a slow rate and in a non-uniform manner that leads to deformed septum shapes if the treatment is performed sufficiently late during constriction (Proctor, Minc, Boudaoud, & Chang, 2012; Ramos et al., 2019; Thiyagarajan et al., 2015). Similarly, mutations in cell wall synthesis proteins affect the ring. In mutated cells where levels of the β-glucan synthase Bgs4 are low, the ring detaches from the septum and the contour formed by the leading edge of the septum does not lie in a plane perpendicular to the long axis of the cell (Munoz et al., 2013). In the *cps1-191* strain that carries a mutation in the β-glucan synthase Bgs1 among others (Dundon & Pollard, 2020), rings both in intact cells and in protoplasts formed by digesting the cell wall away constrict very slowly compared with wild type (Ting Gang Chew et al., 2020).

Many questions about septum growth in fission yeast remain unanswered. What sets the septum shapes observed in electron micrographs (Munoz et al., 2013; Osumi, 2012; Ramos et al., 2019)? How does the septum inner edge remain circular and coplanar, given the thousands of presumably uncoordinated glucan synthases growing the septum? What mechanism creates the particular shape of the septum?

Here we studied contractile ring constriction and coupled processes in the cell-wall enclosed fission yeast *S. pombe*, a uniquely well-characterized cytokinetic system. We developed a molecularly explicit computational model that fully integrates the actomyosin contractile ring dynamics and cell wall growth kinetics. Fission yeast is the only model organism for which this is presently feasible due to the wealth of available data regarding the identities, numbers, and organization of the ring components. The amounts of over 25 ring components were measured over time and many were biochemically characterized (Chen, Courtemanche, & Pollard, 2015; Courtemanche, Pollard, & Chen, 2016; Hayakawa et al., 2020; Pollard & Wu, 2010; Stark, Sladewski, Pollard, & Lord, 2010; Takaine, Numata, & Nakano, 2015; Wu & Pollard, 2005).

Building an integrated computational model of constriction-septation, we find the ring and the septum are essential for each other’s stability. Simulated rings were almost circular and well organized due to this interdependence. Inward forces on the mechanosensitive cell wall synthesis machinery from ring tension suppressed septum roughness by accelerating growth at septum valleys and decelerating growth at peaks. Since ring tension pulled components to the leading edge of the septum, its shape profoundly influenced ring organization. Straight contractile ring bridges snapped away from irregular septum edges, impairing ring organization.

Our model rationalizes a number of unexplained experimental observations which are clearly identified as consequences of the interdependence of these two systems. Mutual positive mechanical feedback mechanisms are triggered by mutations. Simulated rings in *myo2-E1* cells exhibited oval shapes and straight bridges unpeeling from the membrane, which were seen in previous experiments (Laplante et al., 2015; Zhou et al., 2015). Misshapen septa and detached bridges were also observed when the cell wall growth process was disrupted, consistent with phenotypes observed experimentally in cells with reduced levels of Bgs4 (Munoz et al., 2013).

## Results

### Integrated model of contractile ring constriction and septation

To understand and quantify the interaction between the ring and the septum, we created a fully three-dimensional mathematical model of the combined ring-septum system and the coupled processes of ring constriction and septation (Figure 2). We describe our model of the ring in the next subsubsection. In the following subsubsections, we describe the coupling of the ring to the three-dimensional surface representing the plasma membrane and the septum, and how the septum is grown.

#### Molecularly detailed simulation of the *S. pombe* cytokinetic ring

The molecularly explicit mathematical model of the cytokinetic ring was tightly constrained by experiment and was developed in our previous study (McDargh et al., 2021) (Figure 2B, Table S1, also see SI Appendix). The ring was built using coarse-grained components, with their biophysical properties, amounts versus constriction time, and organization consistent with previously measured data. For details see Materials and Methods, Appendix, and Table S1.

##### Node-based organization of myosin II and formin

Representations of ring components are consistent with the local structure of rings measured using super resolution microscopy. Formin Cdc12, myosin II Myo2, and other proteins colocalize in membrane-anchored protein complexes called nodes (Laplante, Huang, Tebbs, Bewersdorf, & Pollard, 2016). In simulations, the ring has ~210 nodes at constriction onset. Cdc12 dimers bind nodes upon binding the ring and then nucleate and grow actin filaments. Each Myo2 cluster represents the heads of 8 Myo2 dimers and is an ellipsoid with dimensions 132 × 102 × 102 nm. Centers of Myo2 clusters and formin dimers maintain fixed distances of 94 nm and 44 nm from the membrane respectively.

##### Other components

Actin filaments are represented as jointed chains with a bending modulus respecting the persistence length of ~ 10 μm measured previously (Ott, Magnasco, Simon, & Libchaber, 1993). The unconventional myosin II Myp2 is not observed in nodes. Myp2 displays actin-dependent ring localization (Laplante et al., 2015; Takaine et al., 2015) and lies further away from the membrane than Myo2 (Laplante et al., 2015; McDonald, Lind, Smith, Li, & Gould, 2017), suggesting that it is not membrane anchored. Myp2 may cluster in constricting rings as it forms puncta (Takaine et al., 2015). Our Myp2 clusters are equivalent to 8 dimers, have a radius of 100 nm and are unanchored (Figure 2B). The cluster representation is consistent with previously measured ring organization and tension (McDargh et al., 2021).

##### Forces

Both Myo2 and Myp2 clusters exert binding forces on actin filaments within range, and pulling forces parallel to the filament obeying a force-velocity relation such that the force decreases linearly with the relative velocity between the filament and the respective cluster (Figure 2B). The cluster force is evenly distributed over all bound filaments. We used stall forces per cluster that reproduce our previously measured experimental ring tensions (Table S1). Drag forces from the cytosol resist component motion. The membrane exerts a drag force as well on anchored nodes. The drag coefficient was chosen such that experimental measurements of node velocities are reproduced (Laplante et al., 2016). Excluded volume forces prevent node-node, node-Myp2, Myp2-Myp2, and filament-filament overlap and filament crossings. Ain1 α-actinin dimers dynamically crosslink filament locations in close proximity.

##### Turnover and numbers of ring components

Rates of component binding and unbinding, and actin filament length dynamics are set by demanding simulations reproduce the experimental densities of Myo2, Myp2 and formin and mean actin length versus constriction time (Courtemanche et al., 2016; Wu & Pollard, 2005). The ring has ~210 nodes, ~140 clusters of Myp2, ~220 formin dimers and ~500 μm of actin filaments at constriction onset. Filaments undergo cofilin-mediated severing (Elam, Kang, & De la Cruz, 2013), Figure 2E.

#### Interaction of the leading edge of the septum with the ring

The septum is a three-layered structure, consisting of an innermost primary septum flanked by two layers of secondary septum, Figure 2A (Garcia Cortes et al., 2016). Electron micrographs from cross-sectional cuts along the long axis of the cell reveal an average thickness ~ 200 nm and a tapering profile, although the amount of tapering is highly variable. The ring shape follows the innermost region of the septum. Viewed along the long axis of the cell, the division plane has a central septum hole, with the ring lining the inner leading edge of the septum (Figure 2A).

The septum leading edge interacts with the ring as follows. Nodes and the formin molecules bound to them are constrained to maintain fixed distances from the membrane, consistent with recent super-resolution measurements (Laplante et al., 2016). Hence, their motions are parallel to the local tangent plane as they move over the septum surface due to various forces. Secondly, incoming nodes bind a region of width *d*_bind_ around the septum leading edge (Figure 2D). While membrane delivery during cytokinesis in fission yeast has been studied (N. Wang et al., 2016), the zone of delivery of new ring components to the membrane during constriction is unclear. We find that a tight zone of localization of width *d*_bind_ = 60 nm is required for the ring to remain properly bundled (see Materials and Methods and Figure 2—figure supplement 1).

#### Septum growth rules and interaction of the ring with the septum

The septum is grown by synthesis of β-glucan and α-glucan by thousands of glucan synthesizers distributed on the septum surface (Arasada & Pollard, 2014; Cortes et al., 2012; Goss et al., 2014; Proctor et al., 2012). Of the three layers of the septum, the innermost primary septum grows the fastest and moves the leading edge inward. The two flanking outer layers of the secondary septum are synthesized more slowly and lead to septum thickening away from the leading edge, Figure 2A (Cortes et al., 2007; Cortes et al., 2012). We represent the two growth processes as a region of high growth at the leading edge of size σ_ps_ and rate υ_0_, and a growth rate attenuated by a factor β_ss_ ≪ 1 away from the edge (Figure 2A). Thus, the mean growth rate along the local normal is given by

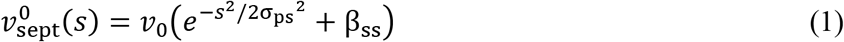

where *s* is the shortest distance of a point on the septum from its leading edge where the ring is located (Figure 2A). We chose values of 60 nm and 0.02 respectively for the size of the high growth region σ_ps_ and growth suppression factor β_ss_ (see subsection “*The contractile ring mechanically regulates septum growth by localization of septum synthesizers*” for details).

Growth rate fluctuations that represent the stochasticity intrinsic to glucan polymerization have zero mean and their correlation is given by

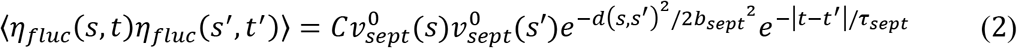

where *b*_sept_ and *τ*_sept_ represent the correlation length and correlation time respectively of septum growth, the normalization constant 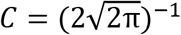, and *d* is the pairwise distance between points(see Materials and Methods for details).

Membrane-bound nodes experience inward normal forces due to being bound to many actin filaments by the Myo2 molecules that they host. In a previous study, we hypothesized that septum growth rates are mechanosensitive and are responsive to inward forces (Thiyagarajan et al., 2015), thus producing a third contribution to septum growth rate in addition to the two terms given by eqns. (1) and (2). Growth rates at a point ***r*** on the septum surface are assumed increased (decreased) when an outward (inward) force from the tension of the ring acts at that point. The change in growth rate is

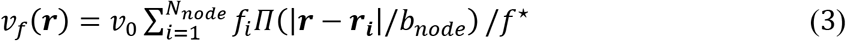

where the positions and inward forces of the *N*_node_ nodes are given by ***r***_*i*_ and *f*_*i*_ respectively, *f*^⋆^ and *b*_node_ represent the critical force at which growth rate is increased and the size of the node respectively, | | represents distances measured along the inner surface of the septum, and Π is the rectangular function of width unity (Figure 2C, see Materials and Methods).

#### Running the simulation

The simulation method is explained in detail in Materials and Methods. The mathematical representation of the septum surface is the set of all points where a function ϕ defined over a three-dimensional volume is equal to zero (Osher & Sethian, 1988). The function was updated at every time step according to the local inward growth rate of the septum. Ring components evolve according to first-order dynamics.

### Contractile rings and septum edges remain almost circular throughout constriction

We simulated ring constriction-septum growth. An initially well-bundled ring of radius 1.85 μm was anchored to an initial septum shape, an annulus with an approximately triangular cross-section whose base width and height were both 200 nm, and equilibrated for 60s after which septum growth was initiated (Figure 3, also see Materials and Methods). The ring maintained an almost perfectly circular contour shape and constricted at a uniform rate, with an almost circular septum hole formed by the septum edge (Figures 3, 4A). The ring tension increased from 530 ± 60 pN at the start of constriction to 1260 ± 180 pN (mean ± s.d., *n* = 4 constrictions) at 17 mins, close to the end of constriction, consistent with our experimental measurements of ring tension (McDargh et al., 2021) (Figure 4B). Cross-sectional profiles of the septum were consistent with previously published electron micrographs (Cortes et al., 2007) (Figure 3, also see the second panel of Figure 4C of the cited reference).

**Figure 3.**
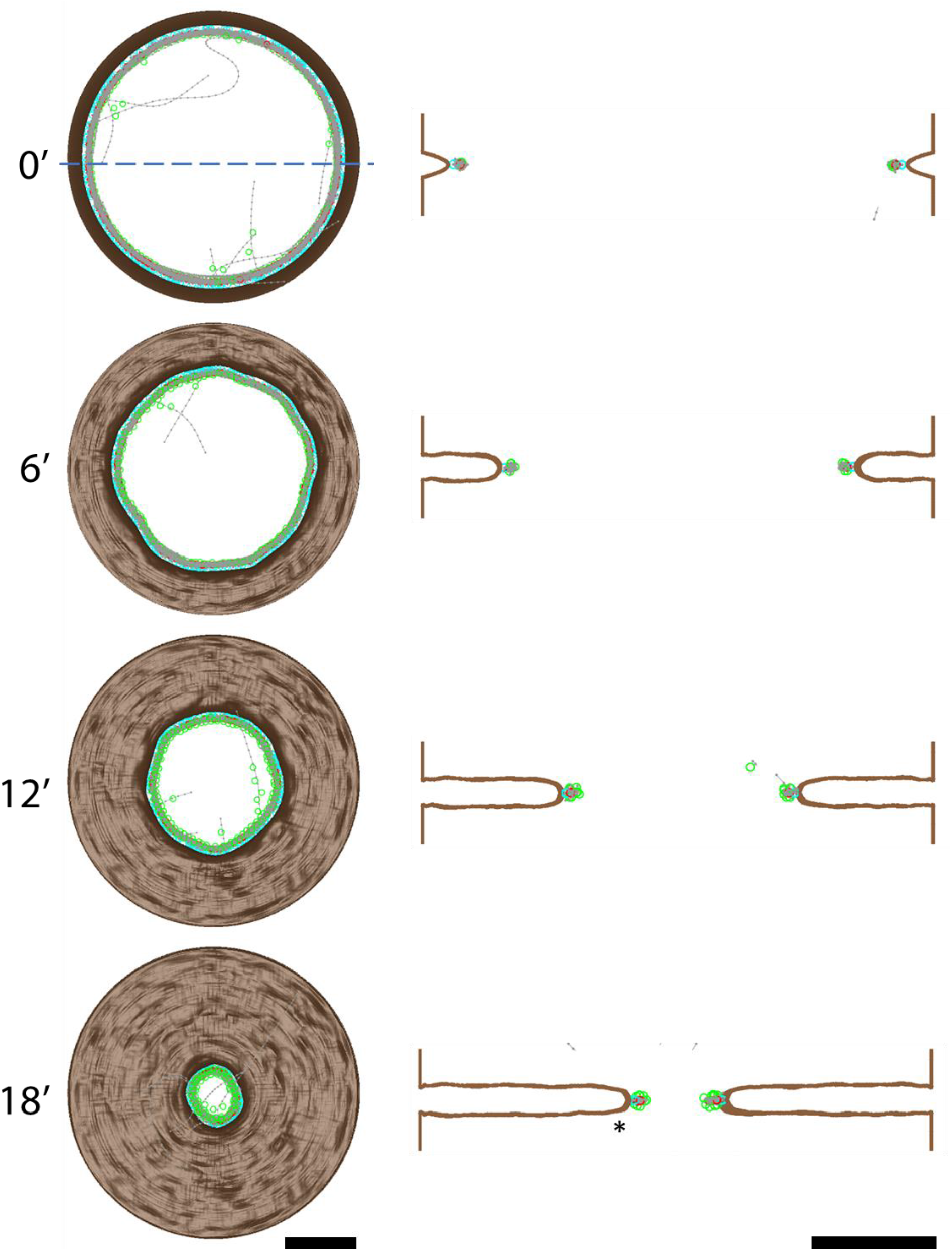
Integrated constriction-septation model produces well-bundled, almost circular constricting rings tethered to a ~200 nm thick septum with almost circular leading edge. Left: Face-on views of a simulated ring constriction-septum growth episode at the indicated times. A well-bundled ring coordinates septum growth, ensuring an almost circular septum hole. Brown: septum surface, red and green circles: Myo2 and Myp2 clusters, grey lines: actin filaments, black lines: actin crosslinkers, which are not prominent as their density is lower than other components. Right: Side views using a section of thickness 200 nm along the blue dashed line of the corresponding ring-septum snapshot at the left. Septa have a mildly tapering profile consistent with experiment, see Figure 6. Asterisk: septum reproduced at greater magnification in Figure 6C. Scale bars are 1μm.

**Figure 4.**
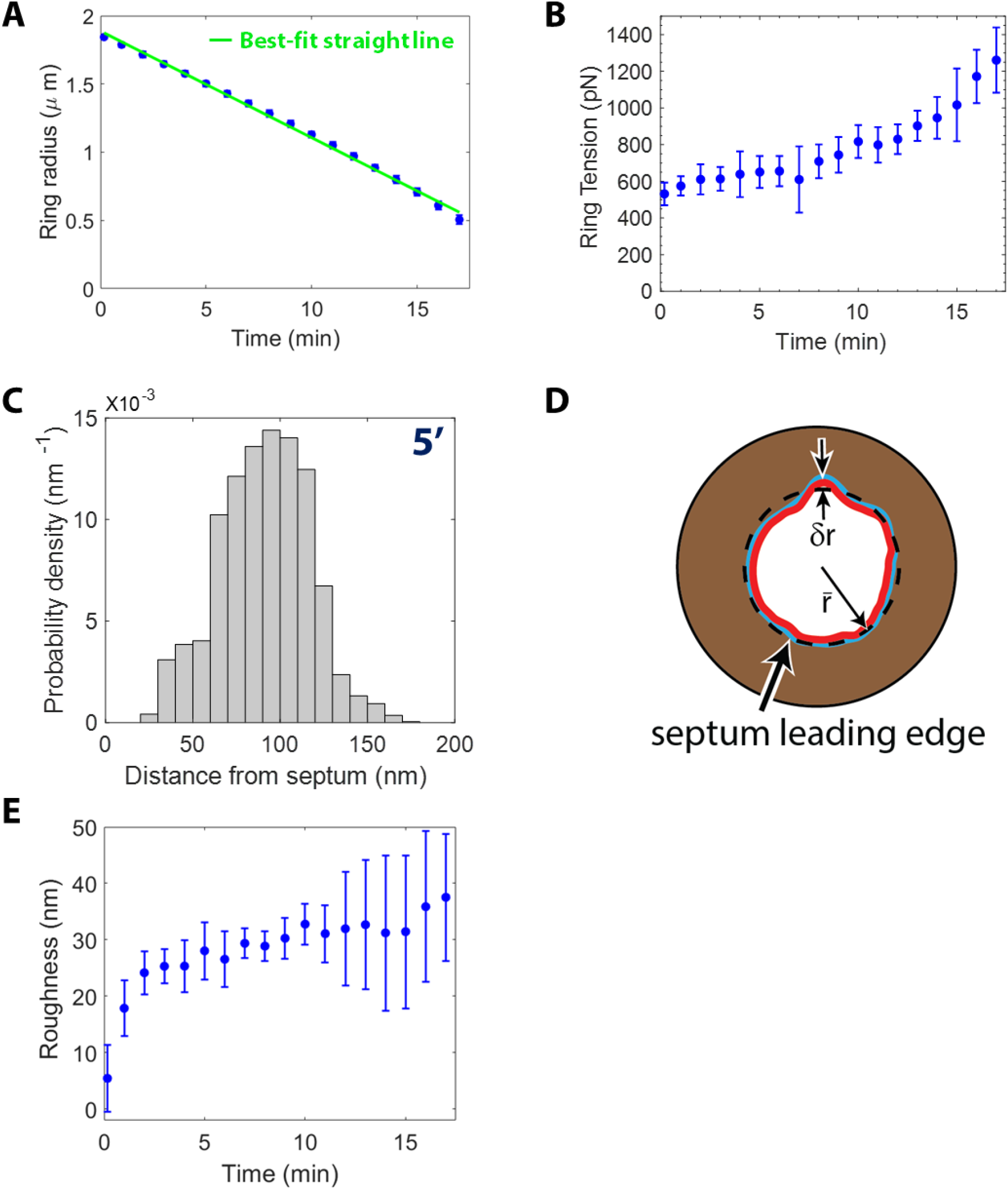
Rings are tense, maintain an almost circular shape and constrict at an approximately constant rate. **(A)** Mean ring radius versus time (mean ± s.d., *n* = 4 constrictions). Rings constrict at an approximately constant rate of 1.28 nm s^−1^. Green: best-fit straight line. **(B)** Mean ring tension versus time for the rings of **(A)** (mean ± s.d.). Ring tension increases from ~500 pN to ~1200 pN over ~16 min. **(C)** Probability distribution function of the shortest distance between actin subunits in a ring and the membrane at the indicated time. A ~30 nm bare zone is present between the ring and the membrane, similar to the ~20 nm bare zone measured using electron cryotomography (Swulius et al., 2018). **(D)** Schematic face-on view of the division plane illustrating roughness measurement procedure. The two-dimensional leading edge of the septum parametrized by *r* is extracted from the top view and its best-fit circle (black dashed line) is obtained whose radius is 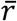. Deviation from circularity 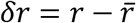 is measured and the roughness 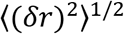 is calculated where 〈 〉 represents spatial averaging over the length of the septum edge contour. **(E)** Time course of roughness of the septum hole of the rings of **(A)** (mean ± s.d.).

The cross-section of the ring 5 min after constriction onset had a thickness 146 ± 13 nm and a width 92 ± 6 nm (mean ± s.d., *n* = 4). There was a gap of size ~30 nm in the direction of the local inward normal to the septum leading edge, where almost no segments of actin filaments were present (Figure 4C). This is similar to the organization observed in (Swulius et al., 2018), where the distribution of distances of ring actin filaments from the membrane showed very few filaments up to a distance ~20 nm from the membrane. These model results are consistent with our previous ring simulations where a similar cross-section was observed when the ring was coupled to a uniformly-ingressing septum represented by the surface of a cylinder (McDargh et al., 2021). After 5 min of constriction, 93 ± 1% of the total actin length and 96 ± 4% of the Myp2 was gathered within the ring (mean ± s.d., *n* = 4). The remaining actin was not bundled and strayed from the ring as “whiskers” decorated sparsely with the remaining Myp2.

We quantified the irregularity of the septum hole seen in the top view of the division plane by roughness *w*. We extracted the septum hole contour parametrized by *r* and measured *w* = 〈(*δr*)^2^〉 where 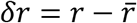 epresents the deviation from circularity of the septum edge, 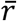 is the mean radius of the septum edge, and 〈 〉 represents spatial averaging over the entire edge (Figure 4D). The roughness plateaus after 5 min into constriction with a slope versus time not significantly different from zero (Figure 4E). The time-averaged roughness measured from 5 min till the end of the simulation was 31 ± 3 nm (mean ± s.d., *n* = 4 constrictions, Figure 4E) consistent with the time-averaged value of 31 ± 9 nm measured using image analysis of septa from live fission yeast cells in our previous study (Thiyagarajan et al., 2015).

### Septum shape regulates contractile ring organization

Next, we examined how ring organization is affected by the septum shape, as the leading edge of the septum provides the backbone to which the ring is tethered. Experiments shows septum shape is crucial for ring viability. In cells of strains where levels of the β-glucan synthase Bgs4 are lower than wild type, the ring detaches from the septum and the contour formed by the leading edge of the septum is tilted and not perpendicular to the long axis of the cell, unlike wildtype (Munoz et al., 2013). Studying growth rate anisotropies is important, as cells of the temperature sensitive *myo2-E1* mutant exhibit anistropic Bgs1 fluorescence around the septum hole and correspondingly anistropic growth rates (Figure 7D of (Zhou et al., 2015)). Thus, abnormal septum shapes due to interference with the cell wall growth process can affect the ring.

We imposed anistropic septum growth to gradually produce a ~1 μm wide high-curvature feature, a “valley”, and used this as an initial condition to observe how the ring-septum system adapted (Figure 5A). To achieve this, the local growth rate was multiplied by an amplitude < 1 compared to the rest of the septum hole using a gaussian function of full width at half maximum 740 nm imposed for a period of 8 minutes from the onset of constriction (Figure 5A, see Materials and Methods).

**Figure 5.**
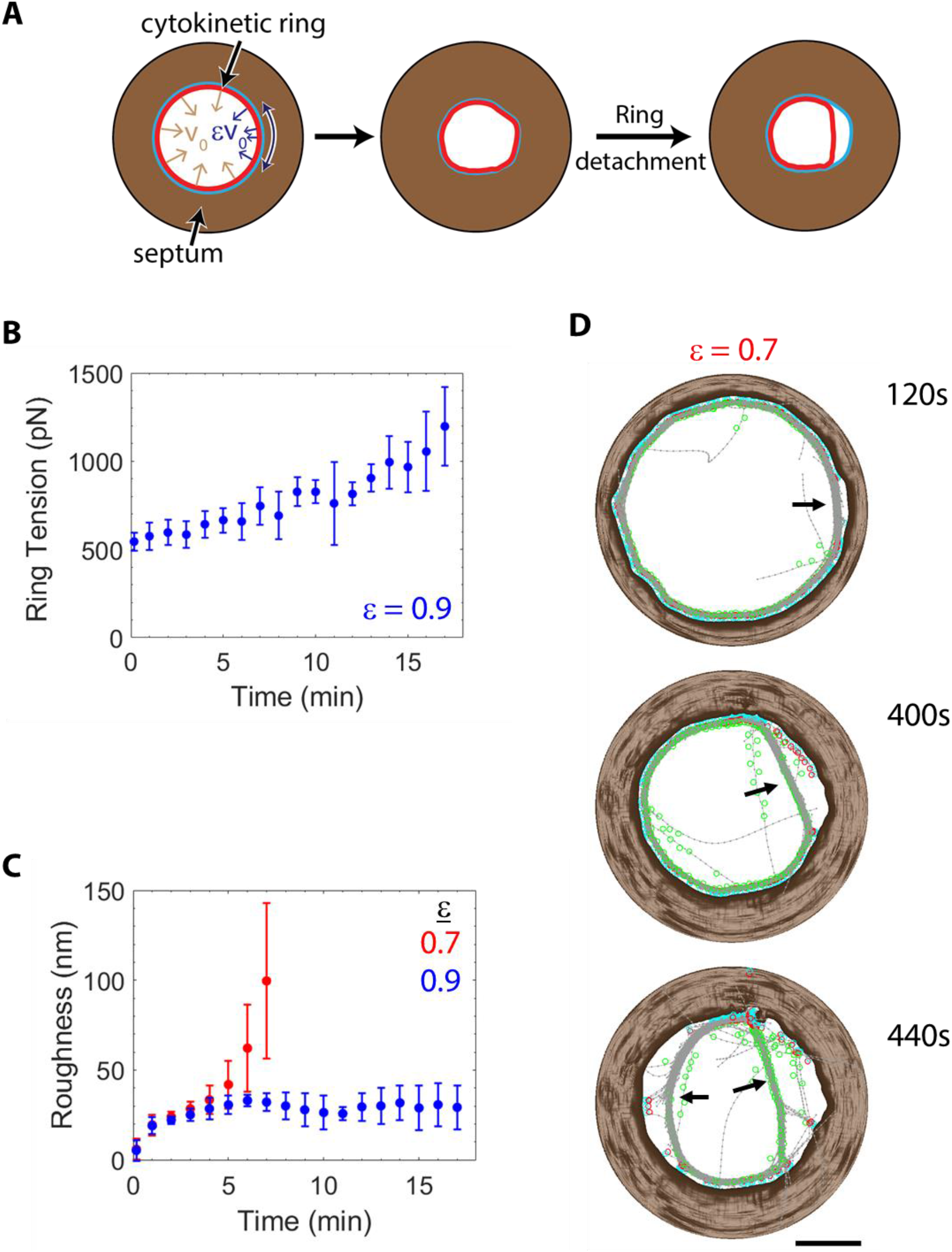
Ring organization is severely disrupted by large septum shape irregularities. **(A)** Schematic showing a local septum growth rate perturbation imposed to examine how septum growth defects affect ring organization. Inward growth rate at a section of the septum was depressed for 8 minutes from constriction onset by a gaussian with amplitude *ɛ* and full width at half maximum of 740 nm, leading to a valley, a region which has lagged in growth (see Materials and Methods). A misshapen septum hole causes ring detachment if its shape is too far from circular. Schematic is not to scale. **(B)** Time course of ring tension for septa with local septum growth rate perturbed to 90% of normal (*n* = 4 constrictions, mean ± s.d.). Tensions are similar to normal simulations with no disruption. **(C)** Time course of septum edge roughness for septa with a local septum growth perturbed to 90% and 70% of normal (*n* = 4 constrictions, mean ± s.d.). Septum shapes are significantly rougher after severe disruptions. **(D)** Time course of face-on view of the division plane of a ring whose local septum growth rate was perturbed to 70% of wild type. A bridge of actin filaments peeled off from the septum at the site of growth perturbation 120 s into constriction (black arrow). The bridge persisted and grew in size. Subsequently another bridge appeared. Scale bar: 1 μm.

We tested different values of the local septum growth rate in the “valley”, from 10% – 90% of normal in steps of 20%. For mild disruption of local growth rates, 90% of normal, rings appeared normal, with roughness and tension similar to wildtype (Figures 5B, C). For 70%, a part of the ring at the “valley” detached from the membrane 1-2 min after the onset of constriction (*n* = 4 rings). The detached part of the ring formed a thick actomyosin bridge, leaving behind a bare zone on the septum (Figure 5D), similar to ring-detachment events observed in cells depleted of Bgs4 (Munoz et al., 2013), and similar to Myp2-decorated bridges that peel off from the membrane in *myo2-E1* cells (Laplante et al., 2016). *Myo2-E1* cells exhibit anisotropic septum growth rates (Zhou et al., 2015). As constriction progressed, other parts of the ring progressively peeled away from the membrane, due to which either the original bridge became longer, or another bridge appeared elsewhere in the ring (Figure 5D). Eventually, the ring was heavily disrupted, and more than half of its length was unanchored. The septum hole never regained circularity (Figures 5C, D). All simulated rings with even higher growth rate disruptions (50% – 10%) suffered similar fates.

Thus, we see that a sufficiently strong perturbation in septum shape leads to drastically impacted ring shape and organization. The ring can no longer guide the septum in the regions where bridges have peeled away, which leads to positive feedback – the septum shape worsens without ring guidance, which in turn worsens ring shape. A circular septum hole, or at least one that is close to circular, is needed for a stable, circular ring.

### The contractile ring mechanically regulates septum growth by localization of septum synthesizers

Experiments show that the ring is intimately coupled to septum growth. When the ring was disassembled using treatment with the actin monomer sequestering drug Latrunculin A, septum ingression either halted or was slow and irregular, leading to thicker septa and non-circular, irregular septum holes (Proctor et al., 2012; Ramos et al., 2019; Thiyagarajan et al., 2015). Full-size rings in cells with mutated Cdc15 slid along the long axis of the cell and stopped sliding when ~2000 molecules of the primary septum synthesizer GFP-Bgs1 had accumulated at the ring (Arasada & Pollard, 2014). Thus, the spatial distribution of septum synthesizers depends on the ring.

In our model, septum growth is fastest at the ring, and within a distance ~σ_ps_ from the ring (Figure 2A). This is because the fast-growing inner primary septum is synthesized by Bgs1 which is enriched at regions of the membrane near the ring (G. Cortés et al., 2015; Goss et al., 2014). Away from the edge region, growth rates are suppressed by a factor β_ss_ and correspond to growth of the secondary septum that flanks the primary septum (eqn. (1)). Using an analytical calculation, we expect septa to have a tapering profile with a mean taper angle *θ*_sept_ ≈ 2β_ss_ (Figure 6A, see Appendix for the calculation).

**Figure 6.**
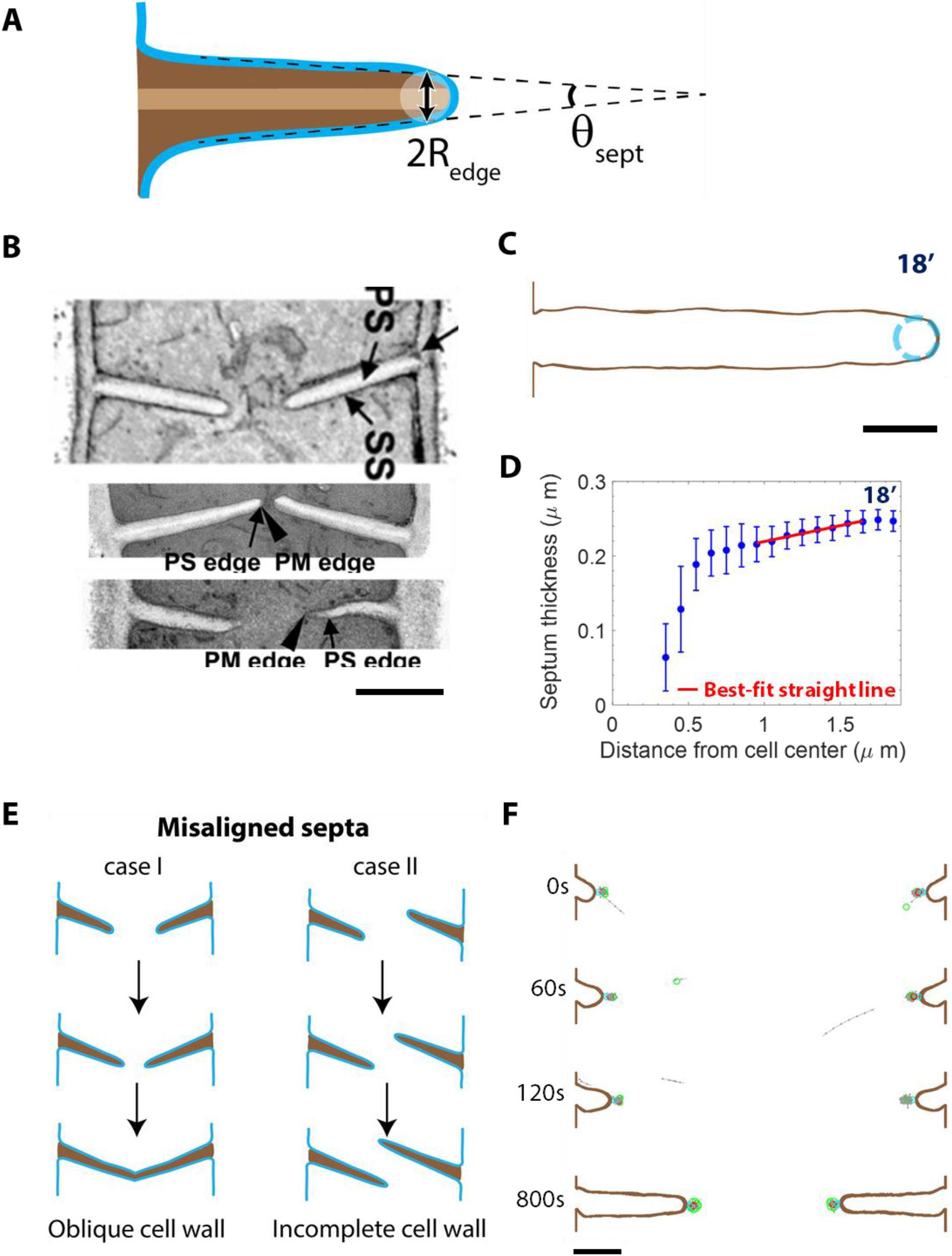
Contractile rings localize septum synthesizers to the septum leading edge, thereby maintaining septum curvature, a tapering septum profile, and suppressing out-of-plane fluctuations of the edge. **(A)** Schematic of septum cross-sectional shape illustrating its geometric characteristics. The radius of curvature of the cross section at the edge is *R*_edge_ and the tapering angle of the cross section is *θ*_sept_, approximately equal to *σ*_ps_ and 2*β*_ss_ respectively (see Appendix for the calculation). **(B)** Electron micrographs of septum cross sections taken along the long axis of the cell. Adapted with permission from Figure 3 of (Munoz et al., 2013) (top) and Figure 9 of (Ramos et al., 2019) (middle and bottom). Septum shapes are tapering and the cross-sectional curvature at the edge varies substantially, with both well-rounded and sharp edges observed. Scale bar: 1 μm. **(C)** Cross-section of the septum of Figure 3 at 18 min (marked ‘*’). Cyan dashed circle: approximate circle fit, radius 86 nm. Scale bar: 300 nm. **(D)** Thickness of the septum versus distance from the central long axis of the cell, averaged over all cross sections of the septum of **(C)** (values are mean ± s.d.). Slope of the best-fit straight line (red) obtained over the window of 0.9 μm to 1.7 μm along the X axis is 0.042 ± 0.028 (68% CI), consistent with twice the value of *β*_ss_ = 0.02 used in the model. **(E)** Schematic of growth with initially misaligned septa. If the growth direction is not corrected, the resultant cell walls are oblique or incomplete. **(F)** Septum and ring cross-sections of a septum with a shape initially misaligned from the division plane by 30^0^ at the indicated times. The septum growth direction became aligned along the plane perpendicular to the long axis of the cell over time, due to ring-mediated localization of septum synthesizers. Scale bar: 500 nm.

In simulations, the ring restricted synthesis to the leading edge and suppressed out-of-plane fluctuations of growth, producing septum cross-sectional shapes similar to electron micrographs published previously (Figures 6B, C). The cross-sectional radius of curvature at the septum edge was ~80 nm, which is similar to the size of the high growth region σ_ps_ = 60 nm (Figure 6C). As these quantities are not different, we see that the ring does not allow primary septum synthesizers to roam freely away from the leading edge. Without such a strong localization effect, we would have observed a radius of curvature much larger than σ_ps_. We also measured a septum taper angle of 2.4^0^ ± 1.6^0^ (68% CI), consistent with twice the value of β_ss_ = 0.02 (Figure 6D).

Our measurements are similar to those we obtained using image analysis of electron micrographs of septa, which validates our parameter choices (Figure 6B). Taper angles varied from ~0.3^0^ to ~3^0^. The radius of curvature of the edge is harder to measure as the inner most point of the septum is unclear. Some edges are rounded, and some appear to be sharp, with a high curvature. We measured the radii of the curvature at previously imaged edges which are relatively well-rounded to be approximately 60 nm (Figure 6B).

The growth of the septum edge must be coordinated in three dimensions to ensure it closes down to a point smoothly. Ring tension pulls synthases to the leading edge, thus setting septum growth direction as normal to the leading edge and ensuring orderly closure.

### The contractile ring corrects an initially misaligned septum by mechanically regulating the localization of septum synthesizers

Electron micrographs from previous studies show that septum edges close to the end of constriction appear to be planar and on course to close smoothly down to a point, Figure 6B (Munoz et al., 2013; Ramos et al., 2019). However, electron micrographs also show septum cross sections earlier during constriction that do not lie in a plane perpendicular to the long axis of the cell (Figure 6B). If such a misaligned septum were to continue its current growth trajectory, the leading edge would not close down to a point, or the resultant daughter cell walls would be oblique as shown in Figure 6E. Course correction of misaligned septa is needed for healthy daughter cells.

To examine if the ring can correct misaligned septum shapes such as those shown in Figure 6E, we simulated ring constriction and septum growth with septum shapes whose cross-sections are initially misaligned by 30^0^ from the plane perpendicular to the long axis of the cell, Figure 6F (see Materials and Methods). The inward growth direction of the septum became perpendicular to the long axis over ~120 s and maintained this orientation for the rest of constriction (Figure 6F). Thus, the ring coordinates septum growth by a unique type of mechanosensitivity—ring tension pulls synthases to the leading edge of the septum and achieves septum coplanarity by suppressing out-of-plane fluctuations. Hence all locations along the edge converge so that the septum hole closes properly.

### The contractile ring regulates septum shape by mechanical regulation of activity of septum synthesizers

Experiments show the contours of the ring and the septum hole are remarkably close to circular throughout constriction (Thiyagarajan et al., 2015; Wollrab, Thiagarajan, Wald, Kruse, & Riveline, 2016; Zhou et al., 2015). A major technical challenge for the cell is to coordinate septum synthesis by thousands of presumably independently operating glucan synthases distributed throughout the septum leading edge (Arasada & Pollard, 2014; Cortes et al., 2012; Goss et al., 2014; Proctor et al., 2012) and maintain circularity.

One part of the septum growth rate in our model is stochastic, with a correlation length *b*_sept_ and correlation time τ_sept_. As synthesis is poorly characterized, we used ring measurements as proxy values. We assumed *b*_sept_ = 150 nm and *τ*_sept_ = 20 s, similar to the thickness of the ring ~125 nm measured using super-resolution microscopy (Laplante et al., 2016) and turnover times of ~10 − 60s of various ring components (Pelham & Chang, 2002; Sladewski et al., 2009; N. Wang et al., 2014; Yonetani et al., 2008). In addition to localizing septum synthesizers to the septum leading edge, the ring also exerts inward pulling forces on the membrane to which it is tethered. We hypothesized that septum synthesis is force sensitive. In our model, growth rate is significantly accelerated compared to the characteristic growth rate υ_0_ at a node occupying a region of size *b*_node_ when it experiences an inward force comparable to a critical force parameter *f*\*.

We simulated constriction episodes with values of *f*\* ranging from 150 to 750 pN. The septum edges were more irregular at higher values of *f*\* (Figure 7A). We quantified the irregularity at the septum edge by roughness, which increased with *f*\* (Figure 7B). A value of *f*\* = 250 pN produced a time-averaged roughness of 30 ± 2 nm (mean ± s.d., *n* = 4 constrictions, Figure 7B) consistent with the time-averaged value of septum roughness of 31 ± 9 nm measured in our previous study using image analysis of septa in live cells (Thiyagarajan et al., 2015). The rate of ring failure also increased with mechanosensitivity *f*\* (Figure 7C), where we defined failed rings as those that bridge and disintegrate before they have constricted down to a radius of 500 nm. The mean septum growth rate decreased mildly with the mechanosensitivity (Figure 7B).

**Figure 7.**
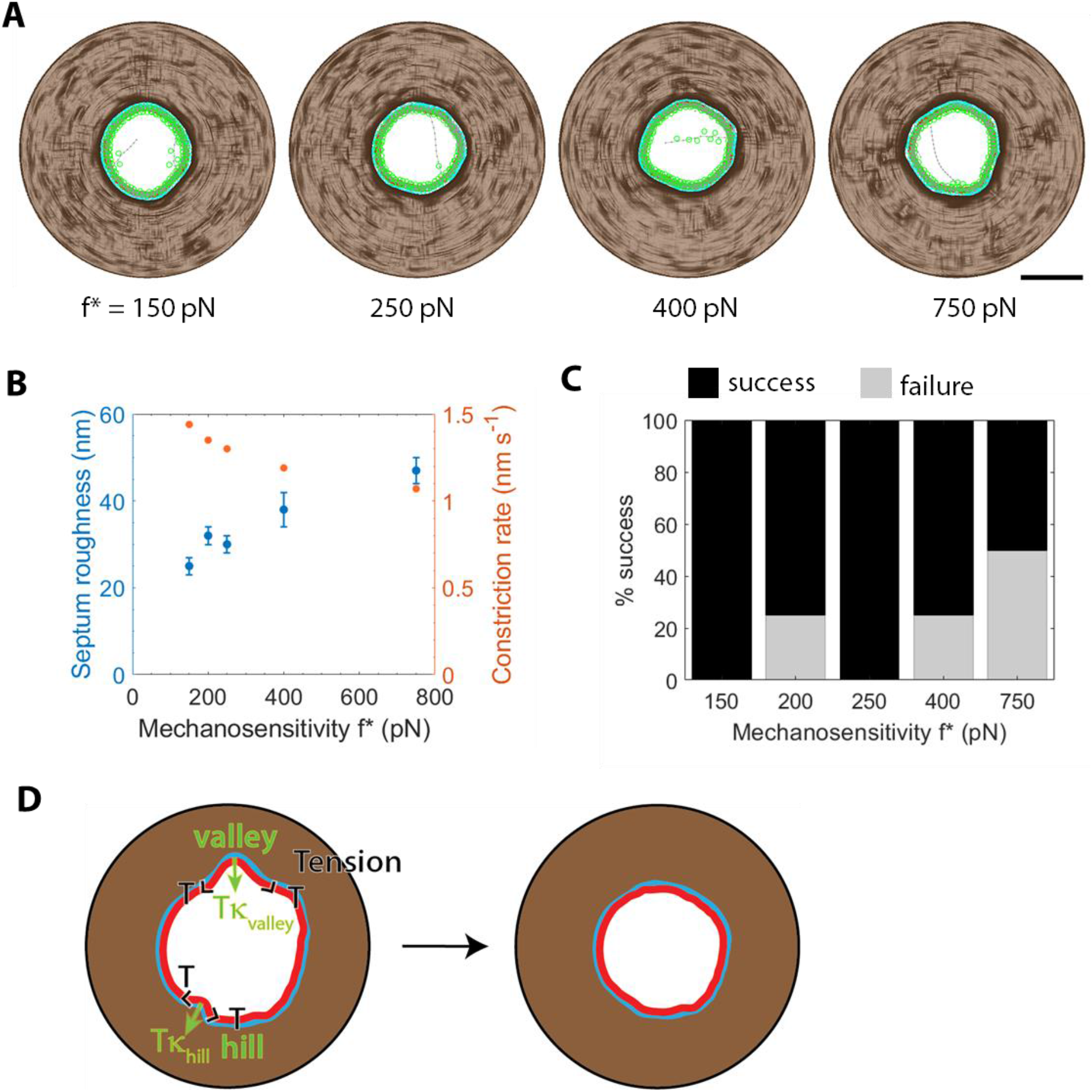
Contractile rings regulate septum growth by exerting forces on mechanosensitive septum synthesizers. **(A)** Face-on view of division plane of ring radius 750 nm at various values of the characteristic force *f*\*, the value of the inward force on a node anchor at which septum synthesis is accelerated by 100% in its neighborhood of size *b*_node_. Septa are rougher at higher *f*\*. The corresponding times after constriction onset are 780 s, 880 s, 960 s, and 1020 s respectively. Scale bar: 1 μm. **(B)** Time-averaged septum roughness and constriction rate versus mechanosensitivity (mean ± s.d., *n* = 4 constrictions each). Roughness was averaged from 5 min to 15 min after constriction onset and constriction rate obtained from the mean ring radius versus time for rings with radii > 400 nm. **(C)** Probability of successful constriction-septation versus mechanosensitivity for the rings of **(B)**. Failure was defined as a ring that bridged and severed at radii > 500 nm. **(D)** Schematic of how ring corrects septum growth through mechanosensitivity of cell wall synthesis. Growth fluctuations can produce a deformed septum edge (left). The inward force exerted by the ring per unit length (arrows) is proportional to local curvature *κ* and ring tension *T* (open arrowheads). Inward force is high at the slow-growing valley leading to local growth acceleration, whereas force is outward at the fast-growing hill leading to local growth deceleration. Thus, the ring compensates for intrinsic fluctuations and ensures almost uniform growth around the septum edge. Circularity is eventually restored (right).

How does the ring suppress septum growth fluctuations? Both in this study and an earlier one that used a much simpler, two-dimensional representation of the septum and an implicit ring (Thiyagarajan et al., 2015), we hypothesize that mechanosensitivity of septum growth is the key. By Laplace’s law, high or low curvature regions experience high or low forces respectively, and growth is adjusted by a factor 1 + *Tκb*_node_/*f*\* where κ is the local curvature of the septum edge (Figure 7D). Thus, growth is decelerated in “hills” and accelerated in “valleys”, which tends to restore circularity of the septum edge.

### Mutations in one component of the ring-septum system destabilize the other component by destructive positive feedback

Although only one experimental measurement probes how septum synthesis mutations affect ring organization to the best of our knowledge (Munoz et al., 2013), many ring mutations that affect septum synthesis have been observed. Septum holes in *cdc12-112* (Thiyagarajan et al., 2015; Zhou et al., 2015), *41xnmtcdc15* (Thiyagarajan et al., 2015), *myo2-E1* (Zhou et al., 2015), and *Rng2pΔIQ* (Thiyagarajan et al., 2015) cells were much rougher than wild-type. We focus in this section on the temperature sensitive *myo2-E1* mutation (Balasubramanian et al., 1998), which lacks Myo2 ATPase activity even at the permissive temperature (Lord & Pollard, 2004).

Curiously, the *myo2-E1* ring and septum phenotypes are variable, a signature of instabilities. The rings appear to be circular (Figure 4D of (Laplante et al., 2016)) or close to circular (Figure S6B of (Zhou et al., 2015)). Straight segments decorated with Myp2 seem to separate from and bridge the ring (Laplante et al., 2016), but are not present in some cells as evidenced by a lack of Rlc1 fluorescence away from the ring contour (Zhou et al., 2015). The septum hole could be oval and off-center (Figure 7D of (Zhou et al., 2015)), close to circular and off-center, or highly irregular with significantly long straight edges (both in Figure 1C of (Zhou et al., 2015)). The size of colonies of *myo2-E1* cells on YES medium is smaller than wild-type cells even at permissive temperatures (Wu, Bahler, & Pringle, 2001), so it is possible that some of these phenotypes are associated with cells that will not survive. Ring tension in *myo2-E1* cells has been experimentally measured to be ~44% of wildtype (McDargh et al., 2021).

We implemented the *myo2-E1* mutation by setting the pulling force per head of Myo2 to zero and decreasing the capture force exerted on actin to 50% of wild-type values. Of the six simulated rings, two developed bridges ~5 min into constriction (Figure 8A). In one ring, multiple bridges occurred rapidly, and the ring was severely disrupted, whereas bridging was transient in the other. The other four rings constricted, albeit with a tension and roughness lower and higher than wildtype, respectively (Figures 4B, 4E, 8B, 8C). At 10 min, the tension was 48% of wild-type, consistent with the ~44% measured experimentally (McDargh et al., 2021), and the roughness was 130% of wild-type. Septum hole shapes were significantly more oval than wild type as constriction progressed (Figure 3, 8A). The phenotype with bridges we observe is similar to the rings in ref. (Laplante et al., 2015), whereas the phenotype with an oval septum hole is similar to the observations of ref. (Zhou et al., 2015). *Myo2-E1* cells also exhibit anisotropic Bgs localization and growth (Zhou et al., 2015). We saw that anisotropic growth produces bridged rings in an earlier section (Figure 5).

**Figure 8.**
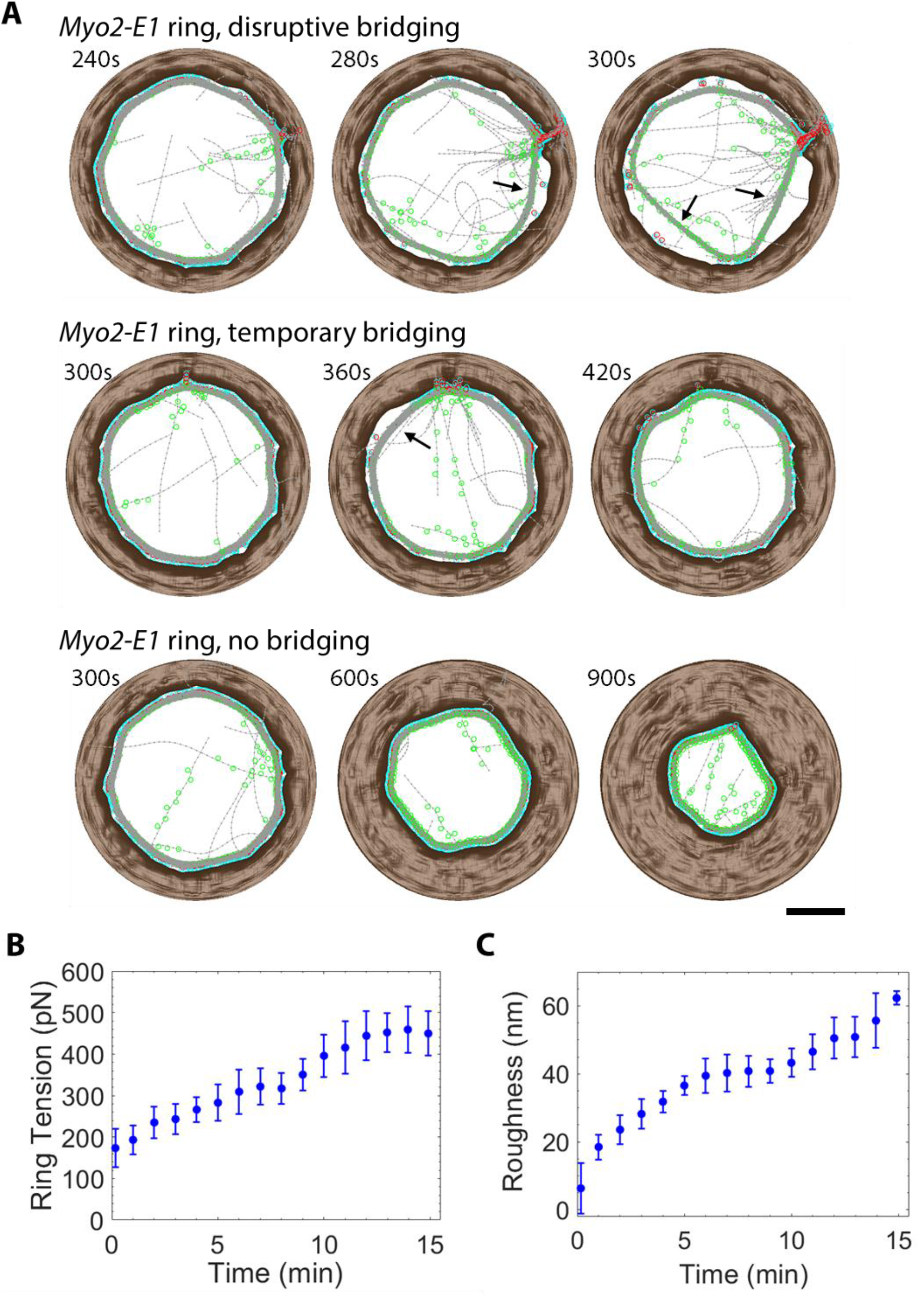
Consistent with experiment, *myo2-E1* rings display two phenotypes: stably constricting rings with oval septum holes, and heavily disrupted rings with multiple bridges. **(A)** Face-on view of the division plane in simulated *myo2-E1* constrictions at the indicated times. Rings show severe bridging (top), transient bridging (middle), or no bridging with oval septum hole shapes (bottom). In the first ring, a thick actomyosin bridge appears 280s into constriction (black arrow). Multiple bridges then appear, disrupting the ring. In the second ring, a short-lived bridge appeared 360s into constriction, and the ring then recovered. In the third ring, constriction appears normal, but septum holes are more oval than wild type. Scale bar: 1 μm. **(B)**, **(C)** Ring tension and septum edge roughness versus time in constrictions with no severe bridging (*n* = 5 rings, mean ± s.d.). Ring tensions are lower than normal simulations, and septa are much rougher.

Thus, we have shown that a sufficiently strong ring perturbation destroys the stabilizing mechanical feedback between the ring and the septum. A highly non-circular ring caused by mutations leads to a non-circular septum. The non-circular septum cannot restore ring circularity and the non-circular ring cannot restore septum circularity. This positive feedback mechanism stabilizes the defective state (Figure 9).

**Figure 9.**
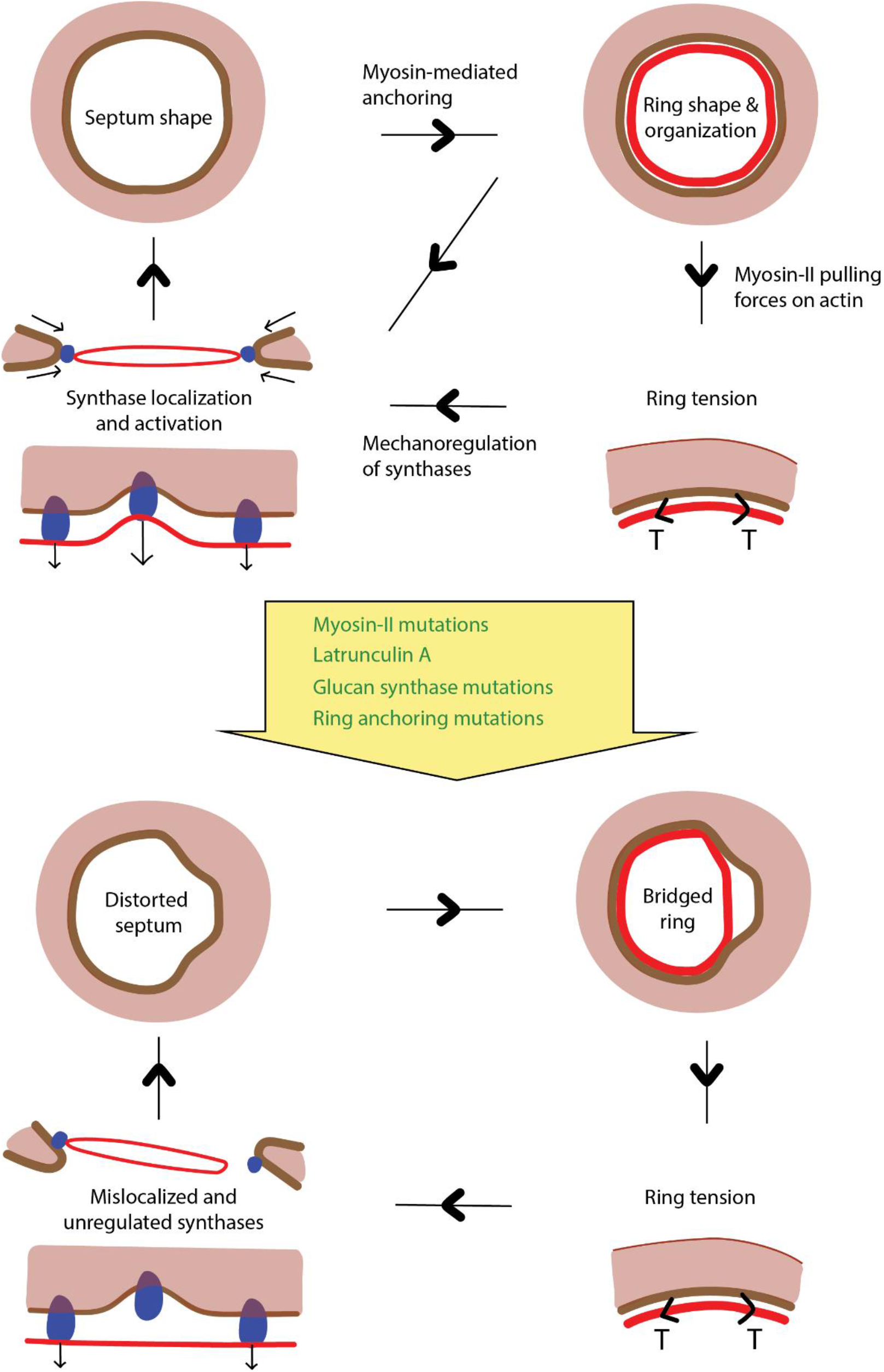
Negative mechanical feedback between ring constriction and septum growth ensures orderly closure, whereas mutations in proteins participating in either process may lead to a runaway positive feedback cascade that disrupts both systems. Top: Schematic of ring constriction-septum growth under normal conditions. The septum furnishes the ring with a circular leading edge, which causes the ring to adopt a circular shape. Ring tension and ring shape perform the two functions of pulling synthases to the leading edge and exerting inward force on them to ensure all regions grow at the same rate. Thus, the synthases maintain a circular leading edge during inward growth, and the feedback loop is complete. Bottom: Schematic of constriction-septation under the influence of one of the listed mutations or drug treatment. As the ring is unable to follow irregular septum shapes, parts of the ring are untethered into bridges. Synthases at the bare regions on the septum do not communicate with the ring and are not regulated as a result. Uncoordinated synthases lead to irregular septum shapes – a positive feedback loop that further worsens ring stability and septum shape.

## Discussion

Our results reveal three important principles in the interdependence between contractile ring constriction and septum growth in fission yeast. (1) First, irregularly shaped interfaces are a threat to ring stability. Due to its tension *T*, the ring experiences centripetal forces per unit length ~ *Tκ* according to Laplace’s law, where κ is the local curvature along the septum hole (Figure 6D). These forces tend to unanchor the ring. Thus, portions of the ring become untethered from the membrane when forced to follow high curvature features. In fission yeast, the ring follows the leading edge of the septum (Figure 2A). We find that misshapen septum holes far from circular cause straight actomyosin bridges to peel away from the membrane and disrupt the ring (Figures 5D, 7C, 8A, 9). In summary, normal ring organization and tension production require the ring to be anchored to a regular geometric backbone (Figure 9).

The second and third principles are that (2) the ring guides septum growth by suppressing in-plane growth fluctuations, and (3) the ring guides septum growth by suppressing out-of-plane growth fluctuations. This guidance by the ring requires two kinds of mechanosensing. The septum is a three-layered, thin tapering structure synthesized by thousands of glucan synthesizers on the membrane with stochastic polymerization processes. Their work must be coordinated to ensure daughter cells are furnished with well-shaped cell wall.

The ring maintains septum growth in the plane perpendicular to the long axis of the cell (principle (3)) through a new kind of mechanosensitivity that we have identified—by localizing activity of cell wall growers to the leading septum edge. The ring seeks to minimize its length due to its tension and thus positions itself at the leading edge which is the shortest closed path along the septum surface. Thus, Bgs1 proteins which are known to localize to the ring (Arasada & Pollard, 2014) gather at the leading edge. In this manner, the fast-growing primary septum synthesized by Bgs1 is guided to grow in the division plane, perpendicular to and close to the ring backbone which is coincident with the leading edge. Thus, the ring corrects growth directions of septa that are initially oblique and ensures daughter cell walls are perpendicular to their axes (Figures 6F, 9).

We propose that the ring maintains circularity of septum holes (principle (2)) by exerting forces on mechanosensitive cell wall growers. The force per unit length exerted by the ring is proportional to local curvature (Figure 7D). We hypothesized in an earlier study, which used a two-dimensional model of septum growth with an implicit ring, that septum growth processes are mechanosensitive and are accelerated by inward ring forces (Thiyagarajan et al., 2015). It follows that growth rate is accelerated in valleys of the septum edge which experience high inward forces and is decelerated in hills where forces are outward (Figure 7D). Thus, the ring helps maintain septum hole circularity by ensuring all regions of the septum edge grow at the same rate. We compared roughness of model-predicted septum edge shapes with experiment and found a mechanosensitivity of 250 pN per node for septum growth rate (Figure 7). It will be of great interest if Bgs mechanosensitivity can be independently tested.

In a normal wild-type cell, negative feedback mechanisms operate between the ring and septum, ensuring coordination between constriction and septum growth in accordance with principles (2), (3) mentioned above (Figure 9). Due to mutual stabilization, septum holes are almost circular, septa lie in a plane perpendicular to the long axis and have a tapering cross section (Figures 3, 6), and the ring is tense and almost circular (Figures 3, 4). The threat to the ring from irregular septum shapes is kept at bay (principle (1)). In the present simulations, ring tension and the organization of actin filaments in the cross section were similar to previous measurements (McDargh et al., 2021; Swulius et al., 2018). We measured primary and secondary septum growth parameters such as the size of the region around the leading edge over which primary septum growth is dominant and the ratio of secondary to primary septum growth rate by comparing model results to previously obtained electron micrographs (Figure 6).

Our results explain destabilizing effects of mutations observed experimentally, showing how ring mutations affect septation and glucan synthase mutations affect the ring (Figure 9). When faced with an irregular septum hole, which may arise from synthase mutations that cause growth problems or ring mutations that lower tension or weaken anchoring, a destabilizing positive feedback mechanism takes hold. Sections of ring untether from the leading edge. Untethered sections can no longer guide septum successfully, so natural stochasticity in growth worsens the septum hole shape, untethering the ring further (Figure 9). Untethered sections of ring are seen in mutants with low Bgs4 levels – a synthase perturbation which can produce growth defects and lead to untethering, consistent with our predictions (Munoz et al., 2013). Bridges and elliptical septum hole shapes were observed in *myo2-E1* cells (Laplante et al., 2015; Zhou et al., 2015). Rings of *myo2-E1* cells have lower tensions (McDargh et al., 2021) which leads to higher roughness and initiates a positive feedback cycle, exacerbated by the fact that the anchoring capacity is lower than for wildtype (Figures 8, 9). Misaligned septa are also seen in *Δmyp2* cells which have lower ring tensions in experiments and many untethered, loose actin filaments, as predicted by our earlier simulation study (McDargh et al., 2021; Okada, Wloka, Wu, & Bi, 2019).

Our work supports the idea that cytokinetic rings are not isolated and are tightly coupled to their environment. Previous experiments measured the effect of isolating fission yeast rings from their normal environment using cell ghosts. In ghosts, the ring environment has been strongly perturbed as the cell wall has been digested away and the membrane has been permeabilized. Here, rings constricted at a rate ~30 times the normal rate (Mishra et al., 2013). The proposed explanation was that large parts of the ring were unanchored, and these sections experienced a non-contractile mode of shortening at the load-free velocity of myosin which is much larger than the normal ring constriction rate (S. Wang & O’Shaughnessy, 2019). Thus, rings experience a drastic change in function when removed from their environment.

Using ghosts, experiments have also measured how the isolation of rings from their normal environment affects their stability. While we have identified irregularities in the inner leading septum edge to which the ring is tethered as a threat to stability (principle (1)), other studies propose that an absence of turnover of ring components is also a threat. Clusters of myosin were observed prominently in rings of cell ghosts that did not constrict (T. G. Chew et al., 2017). In another study, using a mathematical model we concluded that myosin molecules aggregated due to an intrinsic contractile instability arising from regions of high myosin density, which are also regions of high contractile stress, drawing in more myosins from their neighboring regions along the ring length (Thiyagarajan et al., 2021). Turnover of ring components randomizes myosin positions and prevents the onset of aggregation.

As cytokinetic rings are highly dynamic, it is not surprising that their stability and function are heavily dependent on coupled processes. They constrict within a short period of time, with the ring being continuously disassembled and rebuilt simultaneously many times over during constriction as components turn over on timescales which are much smaller than the constriction time (see Table 1 of (O’Shaughnessy & Thiyagarajan, 2018)). The characterization of ring interactions with other cytokinetic processes is key to understanding ring function.

## Acknowledgements

This work was supported by National Institute of General Medical Sciences of the National Institutes of Health under award number R01GM086731 to B.O’S. The content is solely the responsibility of the authors and does not necessarily represent the official views of the National Institutes of Health. We acknowledge computing resources from Columbia University’s Shared Research Computing Facility project, which is supported by NIH Research Facility Improvement Grant 1G20RR030893-01, and associated funds from the New York State Empire State Development, Division of Science Technology, and Innovation (NYSTAR) Contract C090171, both awarded April 15, 2010.

## Author contributions

B.O’S. designed the research. B.O’S. and S.T. performed the mathematical modeling with inputs from Z.M. and S.W., S.T. performed the simulations, B.O’S. and S.T. wrote the paper with inputs from Z.M.

## Data and code availability

The source code of simulations and simulation data are available from the corresponding author upon reasonable request.

## Materials and Methods

### Representation and simulation of the ring

We created a three-dimensional, coarse-grained representation of the cytokinetic ring coupled to the septum surface. In this subsection, we describe the model of the ring. In the next subsection, we describe the septum surface representation, how the septum evolves over time, and how it interacts with the ring. The ring is composed of actin filaments, myosin II clusters, formin dimers, and the actin crosslinker α-actinin. Details of component representation, forces, and turnover processes follow, with key parameters and their numerical values listed in Table S1. For further details, see the Methods section of ref. (McDargh et al., 2021), where the ring simulation was first developed, but was coupled to the inner surface of a constricting cylinder.

#### a. Representation of ring components

The ring model has four components – actin filaments, formin dimers, myosin-II clusters, and actin crosslinkers. We represent actin filaments as a string of beads, with one bead per 100 nm of filament length. The bead at the barbed end corresponds to a formin dimer. There are two types of myosin clusters in our simulation, which represent the two isoforms Myo2 and Myp2. Both myosins are represented by one bead per cluster. Myo2 beads and formin beads are anchored to the membrane. Each Myo2 bead is bound to 0-2 formin beads. One unit of membrane-anchored Myo2 and formin is our representation of the membrane-anchored protein complexes called nodes observed using super-resolution microscopy in the fission yeast ring, Figure 2 (Laplante et al., 2016). Myp2 beads are not membrane anchored. Actin crosslinkers are represented as springs that connect actin beads belonging to different filaments.

#### b. Forces

There are nine types of forces in our model of the ring: (1) myosins capture and exert binding forces on nearby actin filaments, (2) myosins exert pulling forces on bound filaments parallel to the filament length and according to a force-velocity relation, (3) Myo2 clusters and formins experience drag forces parallel to the membrane, (4) Myo2 clusters and formins experience anchoring forces perpendicular to the membrane, (5) crosslinkers exert forces between the pairs of actin beads that they bind, (6) excluded volume forces act between components that are too close to each other, (7) actin filament bending forces and constraint forces between neighboring beads along a filament, (8) cytosolic drag forces, and (9) restoring forces pushing components back into the volume bound by the septum surface if they stray out of it.

##### Myosin capture forces

An attractive capture force with a maximum value of 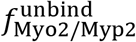 acts between a myosin cluster and an actin bead that has entered the capture zone, which is an ellipsoid with dimensions 132 × 102 × 102 nm and a sphere of radius 100 nm for Myo2 and Myp2 respectively. The Myo2 representation is consistent with super-resolution measurements (Laplante et al., 2016). The force decreases linearly with distance between the bead and the center of the cluster and hits zero at a distance of 25.5 nm for Myo2, which is 1/4^th^ of the length of the minor axis of the Myo2 ellipsoid, and at a distance of 50 nm for Myp2. The force is zero if the captured actin bead is even closer to the center so that filaments have some breathing room and are not all compressed into the center. Only the component of the force perpendicular to the local tangent along the filament length is retained. If there are two beads of one filament within range of a cluster, only the one closest to the pointed end is chosen. If a cluster binds more than 45 filaments (*n*_bound_ > 45), the maximum capture force per cluster-bead interaction is decreased to 45/*n*_bound_ 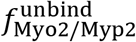.

To suppress numerical instabilities and allow large time steps in the simulation, a pairwise drag force *γ*_a_**Δυ** was added to every pair of actin bead and its bound myosin cluster, where the relative velocity is **Δυ** and the drag coefficient *γ*_a_ = 40 pN s μm^−1^. As this is a pairwise force, no net force was added to the simulation as a result. The force suppresses high-frequency oscillations of the stiff capture springs that are not relevant over long timescales.

##### Myosin pulling forces

Myosin clusters exert pulling forces on bound actin beads parallel to the local tangent to the filament. The force decreases with the relative actin-myosin velocity according to a linear force-velocity relation, with a stall force per filament 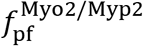 and a load-free velocity 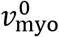. The stall force per filament is given by 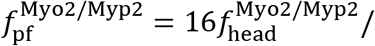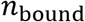, as the pulling force generated by the 16 heads per cluster are divided equally amongst the *n*_bound_ filaments. If the number of bound filaments is too few, *n*_bound_ < *n*_sat_, then the force per filament 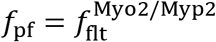, a constant value. It follows that 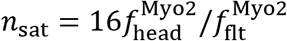.

Our previous study obtained 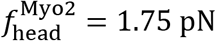 and 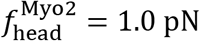 by comparing experimental and simulated time courses of ring tension (McDargh et al., 2021). The study also used 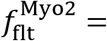 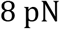. The force per filament was obtained as 4 pN using measured node motions in a previous study of ring assembly (Vavylonis, Wu, Hao, O’Shaughnessy, & Pollard, 2008). It was later ascertained using super-resolution microscopy that one node spot in a confocal image is two nodes that are close together (Laplante et al., 2016). Thus, we revised the per filament value to 8 pN (see (McDargh et al., 2021) for further details). Using these force values, we see that *n*_sat_ = 3.5. We assumed the same *n*_sat_ for Myp2 and obtained 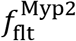 by equating it to 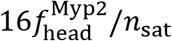.

##### Drag forces from the membrane

A membrane anchored node moves along the membrane and experiences drag forces as a result, with an anchor drag coefficient *γ*_myo_.

##### Anchoring forces

Formin dimers and Myo2 clusters are constrained to lie 44 nm and 94 nm away from the membrane, as reported by super-resolution experiments (Laplante et al., 2016). Anchoring forces enforce the constraint and are calculated implicitly (see (e) of this subsection).

##### Crosslinking forces

Two actin beads from two different filaments which are crosslinked experience an attractive force if they are separated by a distance smaller than 50 nm. The force is linear with a spring constant *k*_x_ and rest length *d*_x_.

##### Excluded volume forces

Ring components separated by small distances experience repulsive forces. The Myp2-Myp2, Myp2-node anchor and Myo2-Myo2 forces are activated if components are separated by distances smaller than 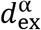 where α can be “Myp2-Myp2”, “Myp2-node” or “Myo2-Myo2”. The forces are linear with respect to the separation between components, with a spring constant 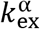 and rest length 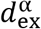(see Table S1). Between actin segments whose minimum separation is *χ*, the excluded volume forces have a mathematical form 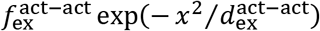.

##### Actin filament bending and constraint forces

Actin filaments represented as semi-flexible polymers. The bead at the barbed end corresponds to a dimer of the formin Cdc12. Constraint forces enforce a separation of 100 nm between neighboring beads along a filament, except the distance between the first two beads whose length varies between 10 nm and 110 nm due to polymerization. When this distance exceeds 110 nm, a new bead is added 10 nm away from the first bead. Bending forces respect the previously measured persistence length of actin (Table S1).

##### Cytosolic drag forces

Actin beads and both myosin clusters experience drag forces with coefficients 0.6 pN s μm^−1^ per bead and 78 pN s μm^−1^ per cluster respectively. We used values higher than what would be expected from viscous interactions with the cytosol as we use the somewhat large time step of 33 ms (see subsubsection (e)) and wanted to ensure numerical stability in the simulations.

##### Confinement forces from the septum

Ring components that are on the wrong side of the septum surface are pushed back with a harmonic restoring force of spring constant *k*_wall_ = 1000 pN μm^−1^.

##### Calculation of ring tension

We calculated the tension of the ring at a point in time by adding the local tensions of all filaments threading one particular cross section and then repeating the procedure for all other cross sections and averaging the measurements.

#### c. Turnover

There are five turnover processes in our model of the ring: (1) Myosin clusters bind and unbind the ring, (2) formins bind new nodes (Myo2 clusters) which are initially devoid of formin, and unbind the ring when the host node unbinds, (3) formins synthesize actin filaments *de novo*, (4) filaments are severed stochastically along their lengths, and (5) actin crosslinkers bind and unbind pairs of filament beads in the ring. The parameters associated with these processes vary with time to respect the experimental measurements of component numbers versus time. Details follow.

##### Myosin turnover

Our model incorporates previous measurements of Myo2 and Myp2 numbers versus time in the ring (Wu & Pollard, 2005), and turnover times of the myosin light chain Rlc1 (Clifford et al., 2008) and Myp2 (Takaine et al., 2015). Myosin clusters are removed stochastically from the simulation according to a mean rate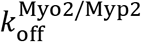 obtained from experiment. As Myo2 and Myp2 numbers vary over time, the on rates 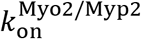 at which new myosins bind the ring are varied in time suitably. Myo2 clusters bind within a distance *d*_bind_ of the leading edge (Figure 2D, also see subsubsection (d) of the next subsection), whereas Myp2 clusters arrive with uniform probability in regions located within 100 nm of actin filament subunits.

##### Formin turnover

In our model, newly arrived nodes are devoid of formin and a node can host a maximum of two formins. Thus, nodes are classified into “zero”, “one”, and “two” nodes depending on the number of bound formins. Formins exit the ring only when their host node is removed. The probability per unit time that a “zero” node is converted into “one”, and a “one” node into “two” are calculated using the equations

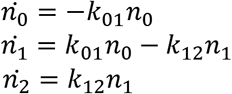

where *n*_0_, *n*_1_ and *n*_2_ are the numbers of the respective node types, and *k*_01_, *k*_12_ are rate constants which do not vary with time. We set 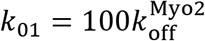 and *k*_12_ = 0.0012*k*_01_. Experimental measurements show that throughout constriction, the ratio of number of formin Cdc12 dimers to number of nodes, counting 16 heads of Myo2 per node, remains close to 1.1 (Courtemanche et al., 2016; Wu & Pollard, 2005). We set the rate constants to respect this ratio, and to ensure the number of “zero” nodes is only ~1% of the number of “one” nodes as we expect formins to bind bare nodes rapidly.

##### Actin turnover

Formins bind nodes and synthesize filaments *de novo* at a rate *υ*_pol_. The ring loses actin stochastically due to two processes: uniform severing at a rate *r*_sev_, which results in the removal of the filament segment from the site of severing to the pointed end, and unbinding of the host node from the ring (see *Myosin turnover* above). The two quantities *υ*_pol_ and *r*_sev_ are time-dependent and are set to reproduce the experimentally-measured time course of mean actin filament length (Courtemanche et al., 2016), and the time over which rings disintegrated upon treatment with a high dose of the actin monomer sequestering drug Latrunculin A (Yonetani et al., 2008). For more details, see ref. (McDargh et al., 2021).

##### Crosslinker turnover

α-actinin on and off rates are set to reproduce numbers and turnover times measured previously (Li et al., 2016; Wu & Pollard, 2005). An incoming α-actinin crosslinker binds a pair of actin subunits selected at random from all pairs belonging to different filaments which are separated by less than 30 nm. In addition to regular turnover, over-stretched crosslinkers with lengths > 50 nm are also removed from the ring.

#### d. Initial ring configuration

The initial ring radius is 1.85 μm. The initial shape of the septum is described in the next subsection. Nodes are initially placed randomly within a narrow band of width 100 nm centered at the septum edge. Each node hosts either 0 or 1 formin dimers, with an average of 0.9 dimers per node. Actin filaments emanating from formins are parallel to the septum leading edge, forming a clockwise or counterclockwise arc with equal probability. Their lengths *l* are drawn from the probability distribution

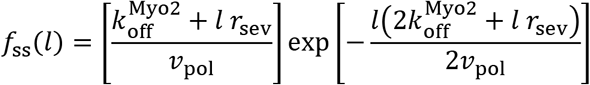

This is the steady-state length distribution of filaments undergoing growth at the rate *υ*_pol_, severing at the rate *r*_sev_ with a uniform probability of severing along their length, and ring unbinding at the rate 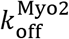. Myp2 is placed randomly within a band that is displaced radially inward by 6 nm compared with the narrow band in which nodes were placed. The ring is equilibrated for 60 s before constriction onset. During the equilibration period, incoming nodes bind within the same narrow band of width 100 nm.

#### e. System of equations and their numerical solution

We describe the equations that are solved to evolve the ring. We used a framework developed previously to enforce constraints (Platt, 1990). Particles whose positions are denoted by {*x*_*i*_} are subject to constraint forces {*λ*_*i*_}, which ensure that the constraints {*g*_*i*_(*x*_*j*_) = 0} are obeyed over a timescale *τ*_c_ = 2/30 s. Constraints enforce anchoring of Myo2 and Cdc12 molecules to the membrane and maintain a distance of 100 nm between consecutive actin beads along a filament (see subsubsection (b) above for more details). Following the Einstein summation convention, the system of equations is:

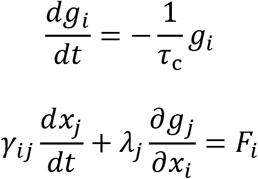

The known variables here are the constraints {*g*_*i*_(*x*_*j*_), the positions of the components {*x*_*i*_}, the forces on components {*F*_*i*_} and the drag matrix *γ*_*ij*_. The interactions described in subsubsection (b) are used to set the forces and the components of the drag matrix, which are set according to the velocity-dependent terms. The equations are solved to obtain the component velocities {*dx*_*i*_/*dt*} and the constraint forces {*λ*_*i*_}. Component positions are then updated according to the forward Euler method. Components are also removed from or added to the simulation according to the turnover rules described in subsubsection (c) above at every time step. We used timesteps of 1/30 s in wild-type simulations and 1/40 s in *myo2-E1* simulations. Results were generated from simulation data saved every 20 s.

### Representation of septum surface and rules of septum growth

#### a. Septum surface representation and evolution

We used the level set method to represent the surface of the septum (Osher & Sethian, 1988). The method represents an (*N* − 1) dimensional surface embedded in an *N* dimensional volume using a scalar function *ϕ*(***r***) defined over the volume. The set of points in the simulation volume that satisfy *ϕ*(***r***) = 0 represent the surface. The value of ϕ at any point in the volume ***r*** is equal to the shortest distance of that point from the surface, which guarantees that 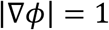.

The surface is evolved over time according to the rule

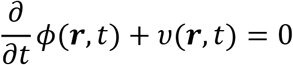

where *υ* is a scalar function with the dimensions of velocity. The local surface growth rate along the local normal is set by the value of *υ* at that point (Osher & Sethian, 1988). We describe *υ* and its components in subsubsection (c) below.

As the surface evolution rule is executed, the nature of *ϕ* as a function that represents distances of points from the surface changes over time. So, over an average of every 120 time steps or 4 s of simulated time, we correct *ϕ* in the neighborhood of the surface using a regularization procedure, which preserves the points at which *ϕ* = 0, and adjusts the values of *ϕ* at other points. We used the distance function of the scikit-fmm package for regularization. We defined *ϕ* within a three-dimensional cuboidal volume with dimensions 4.6 μm, 4.6 μm and 0.6 μm and with a grid size of 10 nm.

#### b. Initial shape of the septum

We chose an initial form for the level set function *ϕ*(***r***) to represent an axisymmetric shape for the septum surface with a tapering cross section. Before regularization, the function is given by

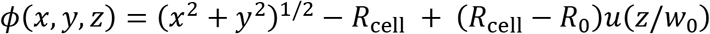

where

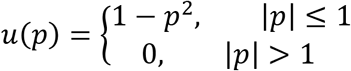

Here, *R*_0_, *R*_cell_ and w_0_ refer to the initial radius of the ring, the radius of the cell membrane, and the initial width of the base of the septum respectively. We chose a tapering form for the function *u* that sets the initial shape of the septum. To ensure *ϕ*(***r***) represents the shortest distance of the point ***r*** from the surface, we regularized it before the rest of the simulation commenced.

For the initially misaligned septum shapes of Figure 6F, we used a cross section whose sides were straight lines oriented at 30^0^ to the inward radial direction. The cross section of the region around the septum edge whose ends were joined with the straight lines was a semicircle with radius 100 nm.

#### c. Rules of septum growth

We evolved the septum surface using the velocity function **υ**(***r***) as described in the previous subsubsection, which is the sum of three components: a mean growth rate 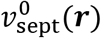, a force-sensitive growth rate **υ**_f_(***r***), and a stochastic growth rate *η*_fluc_(***r***).

### Mean growth rate

We implemented a mean growth rate 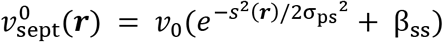 where *s*(***r***) is the shortest distance between the point ***r*** and the ring contour (see the next paragraph for details), and *υ*_0_, β_ss_ (β_ss_ ≪ 1), and σ_ps_ refer to the maximum growth rate, the ratio of the growth rate far away from the leading edge of the septum to the growth rate at the edge, and the width of the region around the edge over which rapid growth occurs.

In practice, it is computationally expensive to calculate the shortest distance *s*(***r***) of all points ***r*** in the simulation volume from the ring contour at every time step. We used an approximation to the gaussian

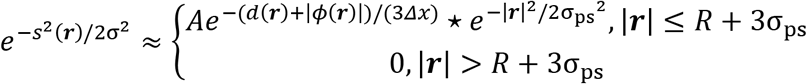

Here ⋆ denotes spatial convolution, *Δx* is the side of one cell in the simulation grid, *R* is the mean ring radius, and *A* is a normalization constant so that the maximum value of the right-hand side is unity. We first calculated the shortest distance *d*(***r***) of every point in space from the best-fit plane to the spatial positions of all nodes. We multiplied the function *e*^−*d*(***r***)/(3Δ*x*)^ with *e*^−|*ϕ*(***r***)|/(3Δ*x*)^ to suppress updating distances at points which are far away from the surface i.e., those that satisfy |*ϕ*(***r***)| ≫ *Δx*. We then convolved the product with a three-dimensional gaussian of width σ_ps_. The resulting function approximates 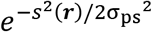, and is highest at the ring and decreases over a length scale σ_ps_ as one moves away from the ring. The mean ring radius *R* is the radius of the best-fit circle to all node positions on the septum.

### Force dependent growth rate

We accelerated septum growth rates at regions which experienced inward force from the ring. The *N*_anc_ nodes anchored at {***r***_*i*_} exert forces {*f*_*i*_} along the local normal, which accelerate growth when they are comparable to a critical force *f*^⋆^. The force-dependent component of the growth rate is 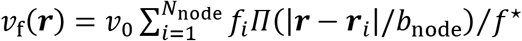, where *b*_node_ represents the width of the region over which the anchor force is felt, and *Π* is the rectangular function of width unity

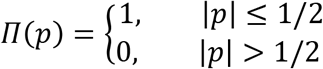

We used time-averaged anchor forces *f*_*i*_ in this rule, obtained by averaging the raw force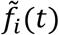 over a moving window of size τ_f_ = 3 s, represented by the equation 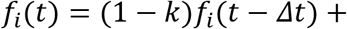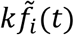, where *k* = *Δt*/τ_f_. Again, to save computational time, we are forced to measure the distances |***r*** − ***r***_*i*_| in three-dimensional space rather than along the surface.

### Stochastic growth rate

We also included a zero-mean stochastic growth rate *η*_fluc_(***r***, *t*) which has a correlation length *b*_sept_ and a correlation time *τ*_sept_ to describe β-glucan synthases operating independently of one another.

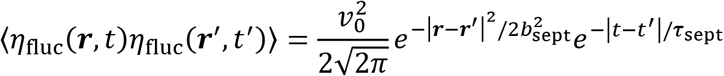

We chose the numerical prefactors 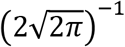 so that at large separations along a straight edge with coordinate *s* and at long times, the statistics of edge roughening produced by the fluctuation is equivalent to that produced by the term

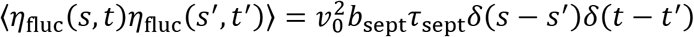

In other words, the prefactors were chosen so that

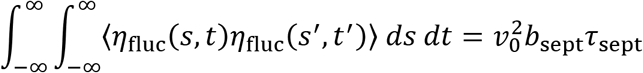

See SI Appendix for further details on the calculation of this integral.

### Local perturbations in growth rate

In the simulations of Figure 5, we introduced a local perturbation only in the mean growth rate component to observe how perturbations in septum shape affect the ring. For 8 minutes from the onset of constriction, we multiplied the mean growth rate by the three-dimensional gaussian function 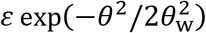 where *θ* is the angular coordinate in the cylindrical coordinate system,*θ*_w_ = 0.17 rad and *ɛ* represent the width and the amplitude of the perturbation respectively. The chosen *θ*_w_ is equivalent to a full-width half-maximum of 740 nm along the leading septum edge at the onset of constriction.

#### d. Binding of incoming nodes

When nodes bind the ring, they do so in regions of high mean growth i.e., in regions on the septum surface where the mean growth component satisfies 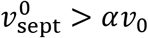 and α is a parameter that sets the cross-sectional width of the binding zone *d*_bind_. Using the exponential functional form of 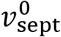 stated in subsubsection (c), we obtain the relation 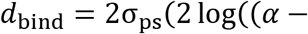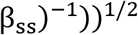 Thus *d*_bind_ = 60 nm corresponds to α = 0.9. To generate results in Figure 2—figure supplement 1, we varied α from 0.1 to 0.9 in steps of 0.2.

