## Supplemental Information Appendix for "Contractile ring constriction and septation in fission yeast are integrated mutually stabilizing processes"

### Analytical calculation of the curvature of the septum tip

We calculate some characteristics of the septum edge shape with a few simplifying assumptions. We assume a cylindrically symmetric septum and ignore fluctuations in growth rate. We assume the ring is perfectly circular and its width and thickness are negligible. Thus, we only consider one cross section of the septum obtained by performing a cut along the long axis of the cell.

As mentioned in the main text, the mean septum growth rate $v_{\text{sept}}^{0}(\boldsymbol{q}) = v_0(e^{-s^2(\boldsymbol{q})/2\sigma_{\text{ps}}{}^2} + \beta_{\text{ss}})$ where $s(\boldsymbol{q})$ is the distance between the point $\boldsymbol{q}$ on the septum surface from the contour of the ring, or equivalently, the leading edge of the septum to a good approximation. Thus, the inward movement of the septum edge and the thickening of the cross section of the septum occur according to the function $v_0(e^{-s/2\sigma_{\text{ps}}{}^2} + \beta_{\text{ss}})$ where $s$ is the arc length coordinate with $s = 0$ representing the leading septum edge (Figure 2A).

We now calculate the cross-sectional curvature $\kappa$ of the septum at the leading edge at steady state. The cross-sectional shape of the septum is represented by the two-dimensional vector function $\boldsymbol{r}(s(t), t)$ where $s(t)$ is the arc length coordinate which is changing in time as the septum grows. The shape evolves according to the equation

$$\frac{\partial}{\partial t}\boldsymbol{r}(s,t) = v_{\text{sept}}^{0}(s)\hat{\boldsymbol{n}}(s)$$

Here, $\hat{n}$ represents the local outward normal. It should be noted that the curvature and the outward normal are related to the septum tip shape as

$$\boldsymbol{r}'' = -\kappa(s)\hat{\boldsymbol{n}}(s)$$

Here $'$ denotes a derivative with respect to $s$. Performing a series expansion of the shape evolution equation around the leading edge $s = 0$, we get

$$\frac{\partial}{\partial t}\left(\boldsymbol{r}(0) + \boldsymbol{r}'(0)s + \boldsymbol{r}''(0)\frac{s^2}{2}\right) = v_0\left(1 + \beta_{\text{ss}} - \frac{s^2}{2\sigma_{\text{ps}}^2}\right)\left(\hat{\boldsymbol{n}}(0) + \hat{\boldsymbol{n}}'(0)s + \hat{\boldsymbol{n}}''(0)\frac{s^2}{2}\right)$$

The terms independent of $s$ cancel, and the various derivatives are as follows. We note that $\boldsymbol{r}'(0)$ and $\boldsymbol{r}''(0)$ are time independent when a steady-state tip shape is achieved.

$$\boldsymbol{r}'(0) = \hat{\boldsymbol{t}}(0)$$
$$\boldsymbol{r}''(0) = -\kappa(0)\hat{\boldsymbol{n}}(0)$$

Here, $\hat{\boldsymbol{t}}(s)$ is the unit tangent vector to the septum tip shape. Using the Frenet-Serret formulas for a plane curve with $\kappa'(0) = 0$, the coefficients on the right-hand side are given as

$$\hat{\boldsymbol{n}}'(0) = \kappa(0)\hat{\boldsymbol{t}}(0)$$
$$\hat{\boldsymbol{n}}''(0) = -\kappa^2(0)\hat{\boldsymbol{n}}(0)$$

Thus, matching the coefficients of the vectors $\hat{\boldsymbol{t}}(0)$ and $\hat{\boldsymbol{n}}(0)$ to second order in $s$ in the shape evolution equation, we get

$$\frac{ds}{dt} = v_0(1 + \beta_{\mathrm{ss}})\kappa(0)s$$

$$-\kappa(0)\hat{\boldsymbol{n}}(0)s\frac{ds}{dt} = -v_0(1 + \beta_{\mathrm{ss}})\hat{\boldsymbol{n}}(0)\frac{s^2}{2\sigma_{\mathrm{ps}}^2} - v_0(1 + \beta_{\mathrm{ss}})\kappa^2(0)\hat{\boldsymbol{n}}(0)\frac{s^2}{2}$$

Using the equations above, we arrive at the result that the curvature of the septum at the tip $\kappa(0)$ is $\sigma_{\mathrm{ps}}^{-1}$.

### Analytical calculation of the taper angle of the septum cross section

A simple geometric argument shows the mean taper angle of the septum cross section $\theta_{\text{sept}}$ is equal to $2\beta_{\text{ss}}$ (Figure 6A). As in the previous section, we parametrize the septum cross-sectional shape using the two-dimensional vector function $\boldsymbol{r}(s(t), t)$. We measure $r$ from the base of the septum. Far away from the leading edge, $\boldsymbol{r}(s,t) \approx (R_{\text{cell}} - R)\hat{\boldsymbol{e}}_{\mathbf{r}} + s\hat{\boldsymbol{e}}_{\mathbf{t}}(s) + \hat{\boldsymbol{e}}_{\mathbf{e}}$, where $\hat{\boldsymbol{e}}_{\mathbf{r}}$ is the inward radial direction, $\hat{\boldsymbol{e}}_{\mathbf{t}}$ is the local tangent, $\hat{\boldsymbol{e}}_{\mathbf{e}}$ is a constant vector set by the cross-sectional shape of the septum near the edge, $R_{\text{cell}}$ is the radius of the cell, and $R$ is the radius of the ring. At steady state, the cross-sectional shape does not change. Differentiating $\boldsymbol{r}(s,t)$ with respect to time at steady state, we get

$$\frac{\partial}{\partial t}\boldsymbol{r}(s,t) = v_0\hat{\boldsymbol{e}}_{\mathbf{r}} + \hat{\boldsymbol{e}}_{\mathbf{t}}\frac{\partial s}{\partial t}$$

We have used the relation that $-\partial R/\partial t = v_0$ and that shape-related vectors do not evolve with time at steady state. Along the sides of the septum, the growth rate is given by

$$\frac{\partial}{\partial t}\boldsymbol{r}(s,t) = \beta_{\text{ss}} v_0 \hat{\boldsymbol{e}}_{\mathbf{n}}$$

where $\hat{\boldsymbol{e}}_{\mathbf{n}}$ is the local normal. Combining both equations and taking a dot product with $\hat{\boldsymbol{e}}_{\mathbf{n}}$ on both sides, we get

$$v_0(\hat{\boldsymbol{e}}_{\mathbf{r}}.\hat{\boldsymbol{e}}_{\mathbf{n}}) = \beta_{\text{ss}} v_0$$

Now, using $\hat{\boldsymbol{e}}_{\mathbf{r}}.\hat{\boldsymbol{e}}_{\mathbf{n}} = \sin\theta_{\text{sept}}/2$, we get

$$\theta_{\text{sept}} \approx 2\beta_{\text{ss}}$$

### Generation of spatiotemporally correlated noise

We use the following algorithm to create a spatially and temporally correlated function $\eta_{\mathrm{fluc}}(\boldsymbol{r},t)$. First, we create three-dimensional, zero-mean gaussian noise, that is neither spatially nor temporally correlated $\eta_{\mathrm{g}}(\boldsymbol{r},t)$, and convolve it with a smoothing function of length scale $b_{\mathrm{sept}}$ and time scale $\tau_{\mathrm{sept}}$

$$\eta_{\mathrm{fluc}}(\boldsymbol{r},t) = \int_{-\infty}^{\infty} dy \int d\boldsymbol{q}\ \mathrm{G}\left(\frac{|\boldsymbol{r}-\boldsymbol{q}|}{b_{\mathrm{sept}}}, \frac{t-y}{\tau_{\mathrm{sept}}}\right)\eta_{\mathrm{g}}(\boldsymbol{q},y)$$
$$\langle \eta_{\mathrm{g}}(\boldsymbol{r},t)\eta_{\mathrm{g}}(\boldsymbol{r}',t')\rangle = v_0^2\delta(\boldsymbol{r}-\boldsymbol{r}')\delta(t-t')$$

Thus, to obtain spatially and temporally correlated noise of the mathematical form

$$\langle \eta_{\mathrm{fluc}}(\boldsymbol{r},t)\eta_{\mathrm{fluc}}(\boldsymbol{r}',t')\rangle = \frac{v_0^2}{2\sqrt{2\pi}} e^{-|\boldsymbol{r}-\boldsymbol{r}'|^2/2b_{\mathrm{sept}}^2} e^{-|t-t'|/\tau_{\mathrm{sept}}}$$

the function $G$ has to satisfy the following convolution relation

$$\int d\boldsymbol{q} \int_{-\infty}^{\infty} dy\ \mathrm{G}\left(\frac{|\boldsymbol{r}-\boldsymbol{q}|}{b_{\mathrm{sept}}}, \frac{t-y}{\tau_{\mathrm{sept}}}\right)\mathrm{G}\left(\frac{|\boldsymbol{r}'-\boldsymbol{q}|}{b_{\mathrm{sept}}}, \frac{t'-y}{\tau_{\mathrm{sept}}}\right) = \frac{1}{2\sqrt{2\pi}} e^{-|\boldsymbol{r}-\boldsymbol{r}'|^2/2b_{\mathrm{sept}}^2} e^{-|t-t'|/\tau_{\mathrm{sept}}}$$

We can see that the following function satisfies this requirement

$$\mathrm{G}\left(\frac{|\boldsymbol{r}-\boldsymbol{q}|}{b_{\mathrm{sept}}}, \frac{t-y}{\tau_{\mathrm{sept}}}\right) = \frac{1}{\pi}\left(\frac{2}{\tau b_{\mathrm{sept}}^3}\right)^{1/2} e^{-|\boldsymbol{r}-\boldsymbol{q}|^2/b_{\mathrm{sept}}^2} e^{-(t-y)/\tau_{\mathrm{sept}}}\theta(t-y)$$

where $\theta$ is the Heaviside step function, which is equal to one for non-negative arguments, and equal to zero everywhere else. We have used the observation that a Gaussian convolved with itself produces a broader gaussian as shown below, where $\star$ represents spatial convolution.

$$\left(\frac{1}{\sqrt{\pi}b_{\mathrm{sept}}}\right)^3 e^{-|\boldsymbol{r}|^2/b_{\mathrm{sept}}^2} \star \left(\frac{1}{\sqrt{\pi}b_{\mathrm{sept}}}\right)^3 e^{-|\boldsymbol{r}|^2/b_{\mathrm{sept}}^2} = \left(\frac{1}{\sqrt{2\pi}b_{\mathrm{sept}}}\right)^3 e^{-|\boldsymbol{r}|^2/2b_{\mathrm{sept}}^2}$$

We chose a gaussian functional form for the spatial correlation as it is easy to obtain the spatial part of the function $G$ analytically. We chose an exponential functional form for the temporal correlation as convolution of a function $f_1$ with $\tau_{\mathrm{sept}}^{-1/2} e^{-t/\tau_{\mathrm{sept}}}\theta(t)$ is equivalent to solving a differential equation in time. In other words, if $f_1^{\tau_{\mathrm{sept}}}$ is the result of smoothing $f_1$ over a time scale $\tau_{\mathrm{sept}}$, $f_1^{\tau_{\mathrm{sept}}} \equiv f_1 \star \tau_{\mathrm{sept}}^{-1/2} e^{-t/\tau_{\mathrm{sept}}}\theta(t)$ where $\star$ represents temporal convolution, then

$$\frac{d}{dt} f_1^{\tau_{\mathrm{sept}}} = -\frac{f_1^{\tau_{\mathrm{sept}}}}{\tau_{\mathrm{sept}}} + \frac{f_1^{\tau_{\mathrm{sept}}}}{\tau_{\mathrm{sept}}^{1/2}}$$

Thus, we need not store and retrieve the entire time series of $f_1^{\tau_{\mathrm{sept}}}$ to perform the convolution, and this allows a faster simulation.

**Supplementary Table 1. Key parameters of the constriction-septation model.**

| Parameter | Meaning | Value | Legend |
|---|---|---|---|
| **Dimensions of the cell and the ring** | | | |
| $R_{\mathrm{cell}}$ | Radius of the cell | 2.06 μm | **(A)** |
| $R_0$ | Ring radius | 1.85 μm* | **(B)** |
| **Actin-myosin capture forces** | | | |
| $f_{\mathrm{Myo2}}^{\mathrm{unbind}}$ | Maximum Myo2 capture force | 40 pN | **(C)** |
| $f_{\mathrm{Myp2}}^{\mathrm{unbind}}$ | Maximum Myp2 capture force | 30 pN | **(C)** |
| **Actin-myosin pulling forces** | | | |
| $f_{\mathrm{Myo2}}$ | Stall force per Myo2 head | 1.75 pN | **(D)** |
| $f_{\mathrm{Myp2}}$ | Stall force per Myp2 head | 1.0 pN | **(D)** |
| $v_{\mathrm{myo}}^{0}$ | Myosin-II load-free velocity | 0.24 μm $s^{-1}$ | **(E)** |
| **Anchoring forces** | | | |
| $\gamma_{\mathrm{myo}}$ | Membrane anchor drag coefficient of a node | 500 pN s $\mu m^{-1}$ | **(F)** |
| **Actin bending forces** | | | |
| $l_{\mathrm{p}}$ | Persistence length of actin filaments | 10 μm | **(G)** |
| **Excluded volume forces** | | | |
| $f_{\mathrm{ex}}^{\mathrm{act-act}}$ | Maximum force of interaction between actin segments | 10 pN | |
| $d_{\mathrm{ex}}^{\mathrm{act-act}}$ | Characteristic range of interaction between actin segments | 15 nm | |
| $k_{\mathrm{ex}}^{\mathrm{Myp2-Myp2}}$ | Spring constant of Myp2-Myp2 interaction | 0.32 pN $nm^{-1}$ | |
| $k_{\mathrm{ex}}^{\mathrm{Myp2-node}}$ | Spring constant of Myp2-node interaction | 0.095 pN $nm^{-1}$ | |
| $k_{\mathrm{ex}}^{\mathrm{Myo2-Myo2}}$ | Spring constant of Myo2-Myo2 interaction | 0.12 pN $nm^{-1}$ | |
| $d_{\mathrm{ex}}^{\mathrm{Myp2-Myp2}}$ | Range of Myp2-Myp2 interaction | 200 nm | |
| $d_{\mathrm{ex}}^{\mathrm{Myp2-node}}$ | Range of Myp2-node interaction | 202 nm | |
| $d_{\mathrm{ex}}^{\mathrm{Myo2-Myo2}}$ | Range of Myo2-Myo2 interaction | 132 nm | |
| **Crosslinking forces** | | | |
| $k_{\mathrm{x}}$ | Spring constant of α-actinin | 25 pN $\mu m^{-1}$ | **(H)** |
| $d_{\mathrm{x}}$ | Rest length of the α-actinin spring | 30 nm | **(I)** |
| **Actin filament growth and severing** | | | |
| $v_{\mathrm{pol}}$ | Actin filament growth rate | 127 nm $s^{-1}$* | **(J)** |
| $r_{\mathrm{sev}}$ | Actin filament severing rate | 0.93 $\mu m^{-1}$ $min^{-1}$* | **(J)** |
| **Myosin turnover** | | | |
| $d_{\mathrm{bind}}$ | Width of region around septum leading edge where incoming nodes bind | 60 nm | **(K)** |
| $k_{\mathrm{on}}^{\mathrm{Myo2}}$ | Myo2 cluster binding rate constant | 0.44 $\mu m^{-1}$ $s^{-1}$* | **(L)** |
| $k_{\mathrm{off}}^{\mathrm{Myo2}}$ | Myo2 cluster unbinding rate constant | 0.0245 $s^{-1}$ | **(M)** |
| $k_{\mathrm{on}}^{\mathrm{Myp2}}$ | Myp2 cluster binding rate constant | 0.31 $\mu m^{-1}$ $s^{-1}$* | **(L)** |
| $k_{\mathrm{off}}^{\mathrm{Myp2}}$ | Myp2 cluster unbinding rate constant | 0.026 $s^{-1}$ | **(N)** |
| **α-actinin turnover** | | | |
| $k_{\mathrm{on}}^{\mathrm{x}}$ | α-actinin on rate | 83 $\mu m^{-1}$ $s^{-1}$ | **(O)** |

| | | | |
|---|---|---|---|
| $k_{\mathrm{off}}^{\mathrm{x}}$ | α-actinin off rate | 3.3 $s^{-1}$ | **(P)** |
| **Septum growth parameters** | | | |
| $v_0$ | Characteristic septum growth rate | 1 nm $s^{-1}$ | **(Q)** |
| $\sigma_{\mathrm{ps}}$ | Width of region around septum leading edge with high growth rate | 60 nm | **(R)** |
| $\beta_{\mathrm{ss}}$ | Ratio of growth rate far away from septum leading edge to growth rate at edge | 0.02 | **(R)** |
| $f^{\star}$ | Critical force of septum growth acceleration | 250 pN | **(S)** |
| $b_{\mathrm{sept}}$ | Correlation length of septum growth | 150 nm | **(S)** |
| $\tau_{\mathrm{sept}}$ | Correlation time of septum growth | 20 s | **(S)** |
| $b_{\mathrm{node}}$ | Size of region on septum surface per node anchor over which mechanosensitive septum growth acceleration occurs | 150 nm | **(T)** |
| $w_0$ | Initial width of the base of the septum | 200 nm | **(U)** |

**Legends**

Note: * signifies that the values are those set at the onset of constriction. Experiments show variables such as the amount of myosin vary with time. Hence, the marked parameters are varied throughout constriction. Also see the corresponding subsections in *Materials and Methods*.

(A) Previous measurements report cell radii lie in the range ~1.5 µm to ~2.2 µm (Facchetti, Knapp, Flor-Parra, Chang, & Howard, 2019).

(B) Largest ring circumferences are in the range ~12-14 µm (Bellingham-Johnstun, Anders, Ravi, Bruinsma, & Laplante, 2021). The chosen ring radius corresponds to a circumference of 11.6 µm.

(C) Obtained in (McDargh et al., 2021) by comparing simulated ring cross-sectional measurements with experiment (Laplante et al., 2015; McDonald, Lind, Smith, Li, & Gould, 2017).

(D) Obtained in (McDargh et al., 2021) by comparing ring tension measured experimentally and in simulations.

(E) Obtained in (McDargh et al., 2021) using measurements from previous *in vitro* gliding filament assays (Stark, Sladewski, Pollard, & Lord, 2010).

(F) Obtained in (McDargh et al., 2021) by comparing simulated node velocities with experimental values (Laplante, Huang, Tebbs, Bewersdorf, & Pollard, 2016).

(G) Measured previously (Ott, Magnasco, Simon, & Libchaber, 1993; Riveline, Wiggins, Goldstein, & Ott, 1997).

(H) Measured in crosslinked actin bundles *in vitro* (Claessens, Bathe, Frey, & Bausch, 2006).

(I) Measured using electron microscopy (Meyer & Aebi, 1990).

(J) Obtained in (McDargh et al., 2021). Also see *Actin turnover* in the *Turnover* subsubsection of Materials and Methods.

(K) Values in the 60-100 nm range lead to well-bundled rings (Figure 2—Figure supplement 1).

(L) Previous experiments report ~2900 and ~1900 heads of Myo2 and Myp2 at the onset of constriction (Wu & Pollard, 2005). Binding constants were inferred assuming a ring circumference of 10 μm in that study, using 16 heads per cluster, and using the off rates mentioned here.

(M) Using Rlc1-GFP FRAP half time of 28 s measured previously (Clifford et al., 2008).

(N) Consistent with Myo3–3mYFP FRAP half time of 32 s measured previously (Takaine, Numata, & Nakano, 2015). Please note that Myp2 is also known as Myo3.

(O) Set to reproduce α-actinin numbers measured in (Wu & Pollard, 2005).

(P) Measured previously in *in vitro* experiments (Li et al., 2016).

(Q) Previous studies report values in the range ~0.7 nm $s^{-1}$ (Bellingham-Johnstun et al., 2021) to ~1.3 nm $s^{-1}$ (Thiyagarajan, Munteanu, Arasada, Pollard, & O'Shaughnessy, 2015) for the rate at which the radius of the ring decreases.

(R) See the subsection *The contractile ring mechanically regulates septum growth by localization of septum synthesizers* of *Results*.

(S) See the subsection *The contractile ring regulates septum shape by mechanical regulation of activity of septum synthesizers* of *Results*.

(T) Similar to ring thickness ~ 125 nm and width of Myo2 distribution in a node ~ 130 nm measured using super-resolution microscopy (Laplante et al., 2016).

(U) Measured approximately from previously published electron micrographs (Munoz et al., 2013; Ramos et al., 2019).

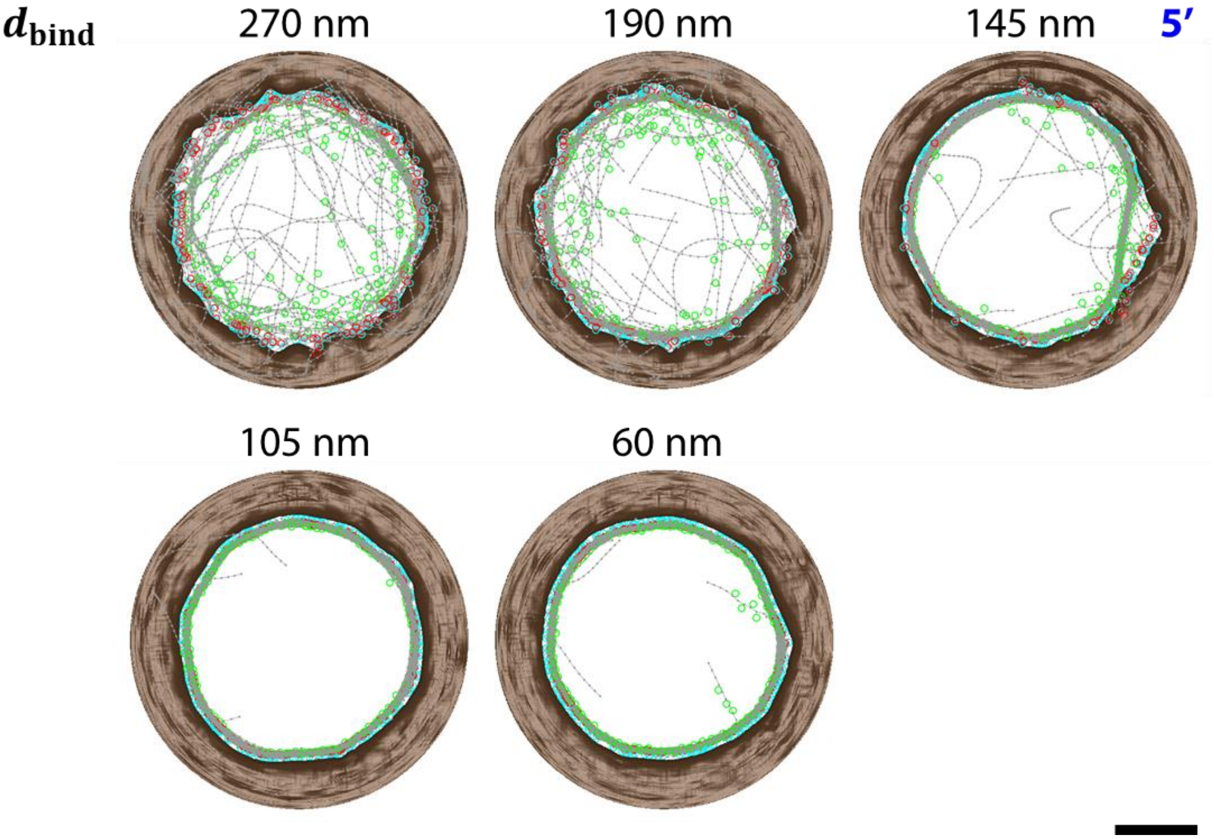


**Figure 2—figure supplement 1**. Top view of simulated division plane at 5 min with the respective width of the binding zone of incoming nodes $d_{\text{bind}}$ (also see Figure 2D). Ring bundling improves as $d_{\text{bind}}$ decreases. Scale bar: 1 µm.
